# Small heat shock proteins HspB1 and HspB5 differentially alter the condensation and aggregation of the TDP-43 low complexity domain

**DOI:** 10.1101/2025.09.17.672772

**Authors:** Thomas B. Walker, Joshua Trowbridge, Shannon McMahon, Nicholas Marzano, Lauren Rice, Justin J. Yerbury, Heath Ecroyd, Luke McAlary

## Abstract

TAR DNA-binding protein 43 (TDP-43) is a nucleic acid-binding protein that regulates processes of mRNA metabolism, during which it undergoes condensation mediated by its C-terminal low complexity domain (TDP-43^LCD^). TDP-43 aggregation and condensation are associated with neurodegenerative disease. However, the proteostasis mechanisms that regulate these processes remain elusive. Some evidence has shown that the molecular chaperone small heat shock protein HspB1 binds to and regulates the cytoplasmic phase separation of TDP-43, indicating that other small heat shock proteins may have similar effects. Here, we demonstrate divergent behaviours for HspB1 and its homolog HspB5 on TDP-43^LCD^ condensation and aggregation. In addition to inhibiting TDP-43^LCD^ aggregation, HspB1 partitions into TDP-43^LCD^ condensates and increases the dynamic exchange of TDP-43^LCD^ within condensates and with the surrounding solution. These effects of HspB1 are enhanced by mutations that mimic phosphorylation. HspB5 inhibits TDP-43^LCD^ aggregation more effectively than HspB1 and partitions into TDP-43^LCD^ condensates, where it delays pathological transition of the condensate to a gel/solid. We localise the chaperone effects of HspB1 and HspB5 to the N- and C-terminal regions of the protein, emphasising the role of sequence diversity in these regions in defining small heat shock protein function. These findings demonstrate that HspB1 and HspB5 are regulators of TDP-43 phase separation and aggregation and may be potential therapeutic targets in mitigating toxic TDP-43 aggregation in neurodegenerative disease.

**Statement:** This work describes how two small heat shock proteins (proteins that bind to misfolded proteins) impact the condensation and aggregation of TDP-43, a protein implicated in most cases of amyotrophic lateral sclerosis. In doing so, it highlights their divergent behaviours and therapeutic potential.

## Introduction

Transactive response (TAR) DNA-binding protein of 43 kDa (TDP-43) is a nucleic acid-binding protein primarily involved in RNA metabolism (1). Normally localised within the nucleus, TDP-43 mislocalises to the cytoplasm of affected cells in amyotrophic lateral sclerosis (ALS) and frontotemporal dementia associated with TDP-43 pathology (FTLD-TDP), where it exists as a major component of pathological proteinaceous inclusions (2, 3). Roughly 22 missense mutations in TDP-43 have been curated as likely pathogenic or pathogenic for ALS and/or FTLD-TDP. Most of these variants are situated within the TDP-43 low-complexity domain (TDP-43^LCD^), a region that governs liquid-liquid phase separation and is aggregation-prone (4, 5), highlighting its potential role in the progression of TDP-43 proteinopathies. Experimental evidence indicates that biomolecular condensates formed via phase separation of TDP-43 may be involved in the pathological mechanisms of aggregation, in which TDP-43 is believed to undergo a phase transition from a dynamic, liquid-like state towards more static, gel-like states (6–8). The mechanisms underlying this process are distinct from classical amyloid fibrillation, in which misfolded proteins rich in β-sheet structures self-assemble into pre-fibrillar oligomers that elongate by irreversibly binding monomers, eventuating in long insoluble fibrils (9). Therefore, to illuminate potential therapeutic strategies against aggregation in ALS and FTLD, it is important to understand the molecular mechanisms that regulate condensation and aggregation of TDP-43.

A core element of protein homeostasis (proteostasis) are molecular chaperones, which are a diverse family of proteins that mitigate protein misfolding. Small heat shock proteins (sHsps) are an important class of molecular chaperones that bind to misfolded proteins and sequester them into large soluble complexes in an ATP-independent manner (10, 11). sHsps are characterised by a central β-sheet-rich α-crystallin domain (ACD), which is flanked by disordered N- and C-terminal regions important for the recognition of client proteins and assembly into large oligomers (12–16). Of the 10 sHsps, HspB1 (also known as Hsp27) and HspB5 (also known as αB-crystallin) are the most widely studied, in part because they have been found to colocalise with astrocytic inclusions in patients with familial ALS (17) and due to their widespread expression and robust chaperone function. These sHsps natively exist as large polydisperse oligomers of up to 40 subunits (11, 18, 19). Phosphorylation of serine residues within the N-terminal domains of HspB1 and HspB5 causes the dissociation of larger oligomers into smaller, more chaperone-active forms, thus enhancing their function during periods of cellular stress (20–25). In line with this, both HspB1 and HspB5 have been found to attenuate aggregation of multiple protein substrates *in vitro* and *in cyto*, a mechanism enhanced by sHsp variants that mimic physiological phosphorylation (24, 26–29).

Despite their importance to proteostasis, the investigation of HspB1 and, to a greater extent, HspB5 as regulators of TDP-43 aggregation has been minimal. Indeed, it was only recently demonstrated that HspB1 regulates the condensation of TDP-43 within the cytoplasm (30). To our knowledge, no such investigations have been undertaken for HspB5 or work to address the impact of these sHsps on the aggregation of TDP-43 in the absence of condensation. To address this gap, we examined the roles of HspB1 and HspB5 in regulating the condensation and aggregation of a purified recombinant form of TDP-43^LCD^. We found that HspB1 promotes the condensation of the TDP-43^LCD^ and enhances molecular exchange of TDP-43 between the condensed and dilute phases. On the other hand, HspB5 does not promote phase separation; rather, HspB5 slowed the time-dependent solidification of TDP-43^LCD^ condensates. Furthermore, we show through kinetic aggregation assays that HspB5 is substantially more effective than HspB1 at preventing the fibrillar aggregation of TDP-43^LCD^, and that the ACD of HspB5 is involved in suppressing fibrillation. Overall, the results indicate that HspB1 and HspB5 both act to prevent TDP-43^LCD^ phase transition and fibrillation; however, they do so to different extents.

## Results

### The TDP-43 low complexity domain (TDP-43^LCD^) forms amyloid fibrils in kinetic aggregation assays

Since the C-terminal LCD of TDP-43 harbours an amyloidogenic core region that is essential for promoting fibrillation (31–33), we first sought to determine the propensity of TDP-43^LCD^ to aggregate using an agitated *in situ* thioflavin-T (ThT) assay. We observed an increase in ThT fluorescence over time with increasing concentrations of TDP-43^LCD^ (**Fig. 1a**). Electron microscopy of samples at the end of this assay revealed the increase in ThT fluorescence was due to the formation of fibrils (**Fig. 1b**). Since there appeared to be concentration-dependent differences in the extent of fibril formation, ThT data were fitted to a Boltzmann sigmoidal curve to derive their elongation rate (i.e. the rate of fibril growth) and time of lag phase (i.e. the time until an increase in fluorescence is observed). We observed a non-monotonic relationship between TDP-43^LCD^ concentration and elongation rate, wherein the elongation rate of TDP-43^LCD^ exhibited a ∼2-fold increase at 10 µM compared to 2.5 µM, while there was only a ∼1.25-fold increase at 50 µM (**Fig. 1c**). We observed that solutions containing the highest concentrations of TDP-43^LCD^ quickly (within minutes of incubation) became turbid (**Fig. S1**), indicating that condensation had likely taken place, thus preventing some monomeric TDP-43^LCD^ from forming fibrils. Despite this, we observed a decrease in the time of lag phase as the concentration of TDP-43^LCD^ increased (**Fig. 1c**). Additionally, we observed a concentration-dependent increase in maximal ThT fluorescence intensity at end point (**Fig. 1d**). These data are consistent with TDP-43^LCD^ undergoing a nucleation-dependent process of fibril formation under these incubation conditions, accordant with the general model by which amyloid fibrils form (31). Having noted the effect of protein concentration and condensation upon the kinetics of TDP-43^LCD^ fibrillation, we performed all subsequent aggregation assays using 10 µM TDP-43^LCD^ to mitigate any confounding effects from its phase separation.

**Figure 1.**
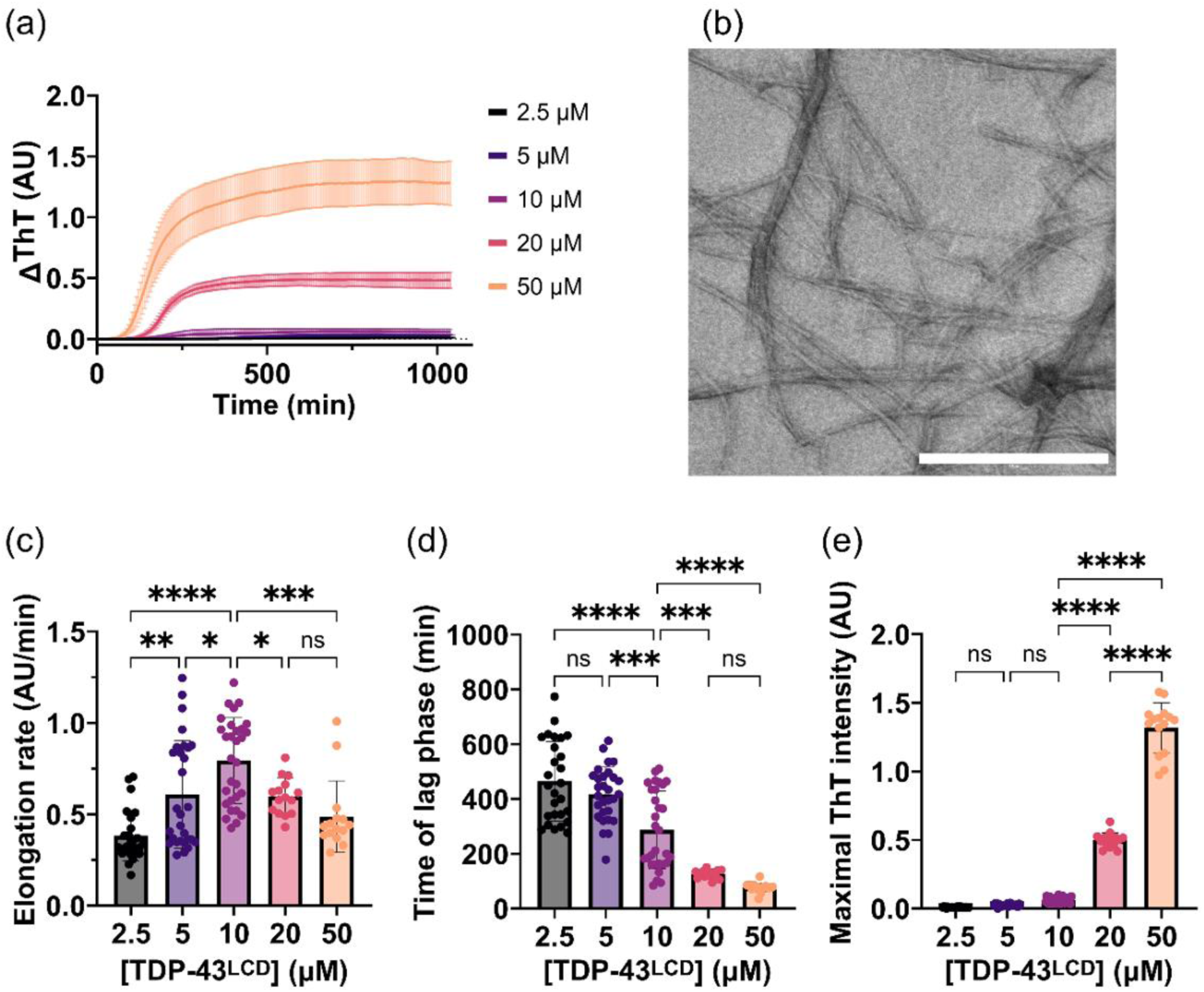
Fibrillation of TDP-43^LCD^ is concentration-dependent. Increasing concentrations of TDP-43^LCD^ (0 – 50 μM) were incubated with 25 μM ThT at 37°C and changes in ThT fluorescence at 490 nm were measured over time. Samples were shaken at 200 rpm for 30 s prior to each cycle. (a) ThT curves for different concentrations of TDP-43^LCD^. (b) Negative stain electron microscope (EM) images of TDP-43^LCD^ fibrils. Scale bar represents 500 nm. (c-e) ThT intensity curves were used to calculate (c) elongation rate; (d) the time of lag phase; and (e) maximal ThT intensity. Data are plotted as mean ± SD (n ≥ 15) and analysed by one-way ANOVA with Tukey’s post-hoc test (* = P < 0.05, ** = P < 0.01, *** = P < 0.001, **** = P < 0.0001).

### HspB5 inhibits TDP-43^LCD^ fibrillation by prolonging the lag phase and slowing elongation to a greater extent than HspB1

HspB1 has been shown to directly bind the LCD of TDP-43 and prevent its assembly into fibrils in cells (30). However, it remains to be established whether HspB1 or HspB5 can alter the kinetics of TDP-43 aggregation in assays using purified recombinant proteins. Therefore, we performed a kinetic ThT assay evaluating the aggregation of TDP-43^LCD^ in the presence of varying concentrations of wild-type forms of HspB1, HspB5 or, as a negative control, bovine serum albumin (BSA). As expected, incubation of TDP-43^LCD^ alone resulted in an increase in ThT fluorescence over time in a sigmoidal manner, indicative of the formation of TDP-43^LCD^ fibrils (**Fig. 2**). Incubation of TDP-43^LCD^ with BSA did not result in any significant change in ThT fluorescence compared to TDP-43^LCD^ alone (**Fig. 2a, b**). However, incubation of TDP-43^LCD^ with HspB1^WT^ or HspB5^WT^ at molar ratios ranging from 5:1 - 1:100 (sHsp:TDP-43^LCD^) inhibited the increase in ThT fluorescence in a concentration-dependent manner (**Fig. 2a, b**). There was no increase in ThT fluorescence when either of the sHsps were incubated in the absence of TDP-43^LCD^.

**Figure 2.**
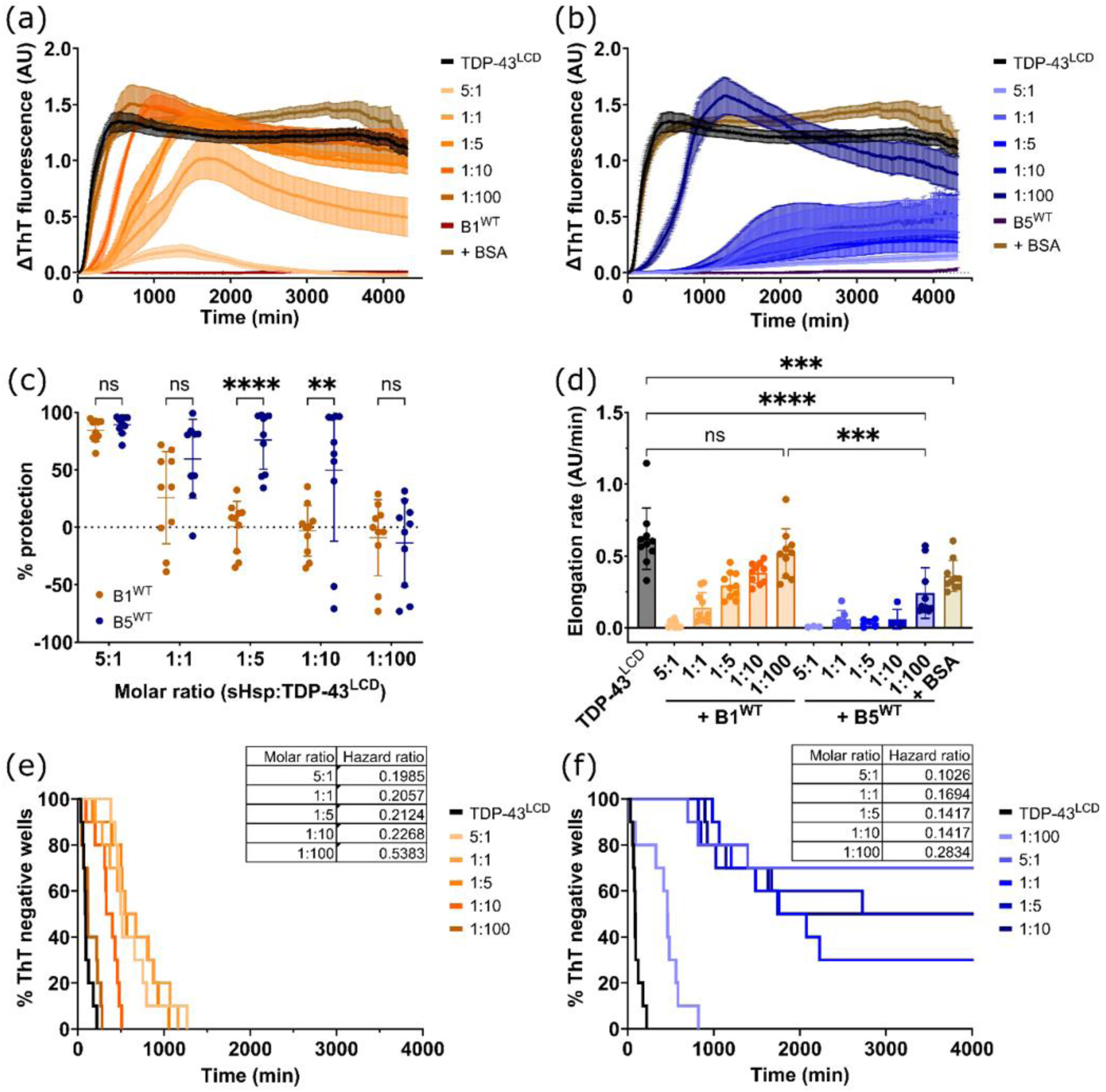
HspB5^WT^ is a more effective inhibitor of TDP-43^LCD^ fibrillation than HspB1^WT^. TDP-43^LCD^ (10 μM) was incubated with HspB1^WT^ or HspB5^WT^ at different molar ratios (5:1, 1:1, 1:5, 1:10, 1:100; sHsp:TDP-43^LCD^). Samples were incubated in the presence of 25 μM ThT at 37°C and monitored for changes in ThT fluorescence at 490 nm. Samples were shaken at 200 rpm for 30 s prior to each cycle. (a and b) ThT curve for TDP-43^LCD^ in the presence of (a) HspB1^WT^ and (b) HspB5^WT^. (c) Percentage protection afforded by HspB1^WT^ and HspB5^WT^ against TDP-43^LCD^ fibrillation. Protection was calculated relative to the mean maximal ThT fluorescence intensity of TDP-43^LCD^ alone. (d) Elongation rate of TDP-43^LCD^ fibrils in the presence of HspB1^WT^, HspB5^WT^ or BSA. (e) Kaplan-Meier analysis of TDP-43^LCD^ fibrillation in the presence of HspB1^WT^. Hazard ratio for each molar ratio is shown inset. (f) Kaplan-Meier analysis of TDP-43^LCD^ fibrillation in the presence of HspB5^WT^. Hazard ratio for each molar ratio is shown inset. Data plotted as mean ± SEM (a, b; n ≥ 9 technical replicates from 2 independent experiments) or mean ± SD (c, d; n ≥ 3 technical replicates from 2 independent experiments). Data analysed by (c) two-way ANOVA with Šidak’s post-hoc test or (d) one-way ANOVA with Tukey’s post-hoc test (** = P < 0.01, *** = P < 0.001, **** = P < 0.0001).

The percentage protection afforded against TDP-43^LCD^ fibrillation by HspB1^WT^ and HspB5^WT^ was similar at molar ratios of 5:1 or 1:1 (sHsp:TDP-43^LCD^) but was greater for HspB5^WT^ at 1:5 and 1:10 molar ratios, suggesting that HspB5^WT^ more potently inhibits the fibrillation of TDP-43^LCD^ (**Fig. 2c**). At lower molar ratios, the presence of the sHsps sometimes resulted in higher levels of ThT fluorescence compared to TDP-43^LCD^ alone (measured as negative percent protection in **Fig. 2c**), indicative of the inability of the chaperones to prevent the overall amount of TDP-43^LCD^ fibril formation at these molar ratios. Fitting to a Boltzmann sigmoidal curve revealed that the presence of the sHsp also led to a decrease in the elongation rate of TDP-43^LCD^ fibrils, indicating that both HspB1^WT^ and HspB5^WT^ impede the growth of TDP-43^LCD^ fibrils (**Fig. 2d**). Even at molar ratios as low as 1:100, HspB5^WT^ was capable of significantly reducing the elongation rate of TDP-43^LCD^ fibrillation compared to TDP-43^LCD^ alone, while HspB1^WT^ had no effect on the elongation rate at this same molar ratio. Whilst BSA significantly reduced the elongation rate of TDP-43^LCD^ fibrils, this only occurred at the relatively high molar ratio of 1:1. Thus, these data indicate that HspB5^WT^ is more effective than HspB1^WT^ at reducing TDP-43^LCD^ fibril formation under these conditions.

A Kaplan-Meier analysis of the lag-phase of aggregation was performed as described previously (34) to assess the impact of the chaperones on the initiation of fibril formation by TDP-43^LCD^ within individual wells of the assay. In the presence of HspB1^WT^, the onset of TDP-43^LCD^ fibrillation was significantly delayed at molar ratios as low as 1:10 (sHsp:TDP-43^LCD^), as indicated by the right-shift in the Kaplan-Meier distributions (**Fig. 2e**). The onset of TDP-43^LCD^ fibrillation was significantly delayed in the presence of HspB5^WT^ at each of the molar ratios tested (**Fig. 2f**). Indeed, HspB5^WT^ consistently conferred a lower hazard ratio than HspB1^WT^ at each molar ratio tested, indicating that it more effectively reduced the likelihood of TDP-43^LCD^ fibrillation (**Table S2**). Altogether, these data indicate that HspB5^WT^ is a potent inhibitor of the formation and growth of TDP-43^LCD^ fibrils.

### A phosphomimetic isoform of HspB1 (HspB1^3D^) is more effective than wild-type HspB1 at preventing TDP-43^LCD^ fibrillation

Previous studies have shown that phosphomimetic mutations at serines 15, 78 and 82 reduce the size of HspB1 oligomers, while similar mutations in HspB5 have a lesser effect upon its oligomeric distribution (21, 24, 35). Furthermore, phosphomimetic HspB1 has been found to have enhanced chaperone activity against several aggregating proteins through increased exposure of substrate binding regions (24, 36). We therefore sought to examine whether HspB1^3D^ is more effective than HspB1^WT^ at preventing the formation of TDP-43^LCD^ fibrils. Incubation of TDP-43^LCD^ with increasing amounts of HspB1^3D^ led to a decrease in ThT fluorescence intensity compared to when TDP-43^LCD^ was incubated alone (**Fig. 3a**). Moreover, the presence of HspB1^3D^ conferred protection against TDP-43^LCD^ fibrillation at all molar ratios tested (**Fig. 3b**). This contrasted with HspB1^WT^, which provided no protection against TDP-43^LCD^ fibrillation at molar ratios of 1:5, 1:10 and 1:100 (**Fig. 2c**). These data are consistent with HspB1^3D^ having greater chaperone activity than HspB1^WT^ (24).

**Figure 3.**
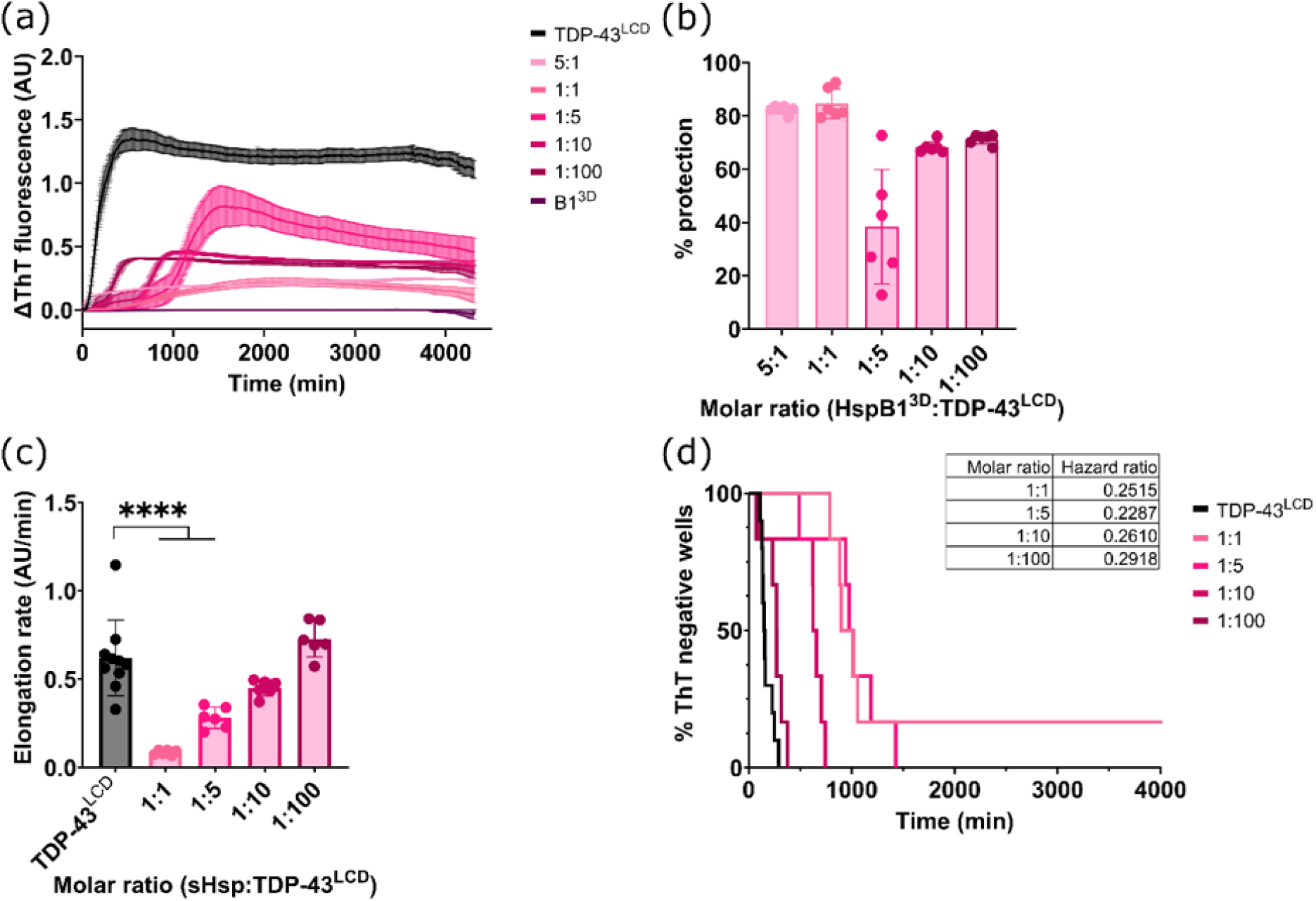
The effect of a HspB1 phosphomimetic on the fibrillation of TDP-43^LCD^. TDP-43^LCD^ (10 μM) was incubated with different molar ratios (5:1, 1:1, 1:5, 1:10, 1:100) of HspB1^3D^ (sHsp:TDP-43^LCD^) and 25 μM ThT at 37°C and monitored for changes in ThT fluorescence at 490 nm. Samples were shaken at 200 rpm for 30 s prior to each cycle. Assay was performed concurrently to assays shown in Fig. 2. (a) ThT curve for TDP-43^LCD^ in the presence of HspB1^3D^. (b) Percentage protection against TDP-43^LCD^ for HspB1^3D^.(c) Elongation rate of TDP-43^LCD^ fibrils in the presence of HspB1^3D^. (d) Time of lag phase for TDP-43^LCD^ fibrillation in the presence of HspB1^3D^. Data plotted as mean ± SEM (a) or mean ± SD (b-d) (n ≥ 5 technical replicates from 2 independent experiments). Data analysed by (c) one-way ANOVA with Tukey’s post-hoc test (**** = P < 0.0001).

The elongation rate of TDP-43^LCD^ fibrils was reduced in the presence of higher concentrations of HspB1^3D^ (i.e. at 1:1 and 1:5 molar ratios sHsp:TDP-43^LCD^), indicating that HspB1^3D^ was impeding the growth of TDP-43^LCD^ fibrils under these conditions (**Fig. 3c**). Additionally, HspB1^3D^ delayed TDP-43^LCD^ aggregation at 1:1, 1:5, 1:10 and 1:100 molar ratios (**Fig. 3d**). Under these conditions, HspB1^3D^ reduced the hazard ratio for TDP-43^LCD^ aggregation, indicating that it was reducing the likelihood of TDP-43^LCD^ aggregation (**Table S2**). Notably, ThT fluorescence data for TDP-43^LCD^ in the presence of HspB1^3D^ at a 5:1 molar ratio could not be fitted to a Boltzmann sigmoidal curve. Altogether, these findings are consistent with previous work demonstrating that phosphomimetic mutant HspB1 exhibits greater chaperone activity than its wild-type counterpart (24, 27).

### The α-crystallin domain (ACDs) of HspB1 and HspB5 partially inhibit TDP-43^LCD^ aggregation

Given that both the full-length isoforms of the sHsps inhibit the aggregation of TDP-43^LCD^, we next determined whether this is mediated by the conserved ACDs of these sHsps. Surprisingly, at a 1:1 molar ratio (sHsp^ACD^:TDP-43^LCD^) HspB1^ACD^ increased the maximum ThT fluorescence compared to when TDP-43^LCD^ was incubated alone (**Fig. 4a**). Importantly, control wells containing HspB1^ACD^ alone showed no increase in ThT signal, indicating that this increase in ThT fluorescence was associated with TDP-43^LCD^ aggregation (**Fig. 4a**). In contrast, levels of ThT fluorescence associated with TDP-43^LCD^ fibril formation were reduced in the presence of HspB5^ACD^ at a 1:1 molar ratio (sHsp^ACD^:TDP-43^LCD)^ (**Fig. 4b**). Thus, in these experiments, HspB5^ACD^ conferred greater protection against TDP-43^LCD^ aggregation than HspB1^ACD^; although, it provided only modest protection at a 1:1 molar ratio and had little to no effect at 1:10 and 1:100 molar ratios (**Fig. 4c**). In addition, whilst HspB1^ACD^ had no effect upon the elongation rate of TDP-43^LCD^ fibrils at any of the molar ratios tested, HspB5^ACD^ reduced the elongation rate at a 1:1 molar ratio (**Fig. 4d**). A Kaplan-Meier analysis demonstrated that TDP-43^LCD^ aggregation was less likely to occur in the presence of an equimolar amount of HspB1^ACD^, which yielded a significantly reduced hazard ratio at this molar ratio; however, no difference was observed at a 1:10 or 1:100 ratio (HspB1^ACD^:TDP-43^LCD^) (**Fig. 4e**). In contrast, TDP-43^LCD^ aggregation was relatively unaffected by HspB5^ACD^ at a 1:1 or 1:10 molar ratio but was accelerated at a 1:100 molar ratio (**Fig. 4f**). Under this condition, HspB5^ACD^ conferred a much greater hazard ratio, indicating that it was increasing the likelihood of TDP-43^LCD^ aggregation (**Table S2**). These findings suggest that the ACDs of these two sHsps play only a relatively minor role in the inhibition of the aggregation of TDP-43^LCD^ into fibrils.

**Figure 4.**
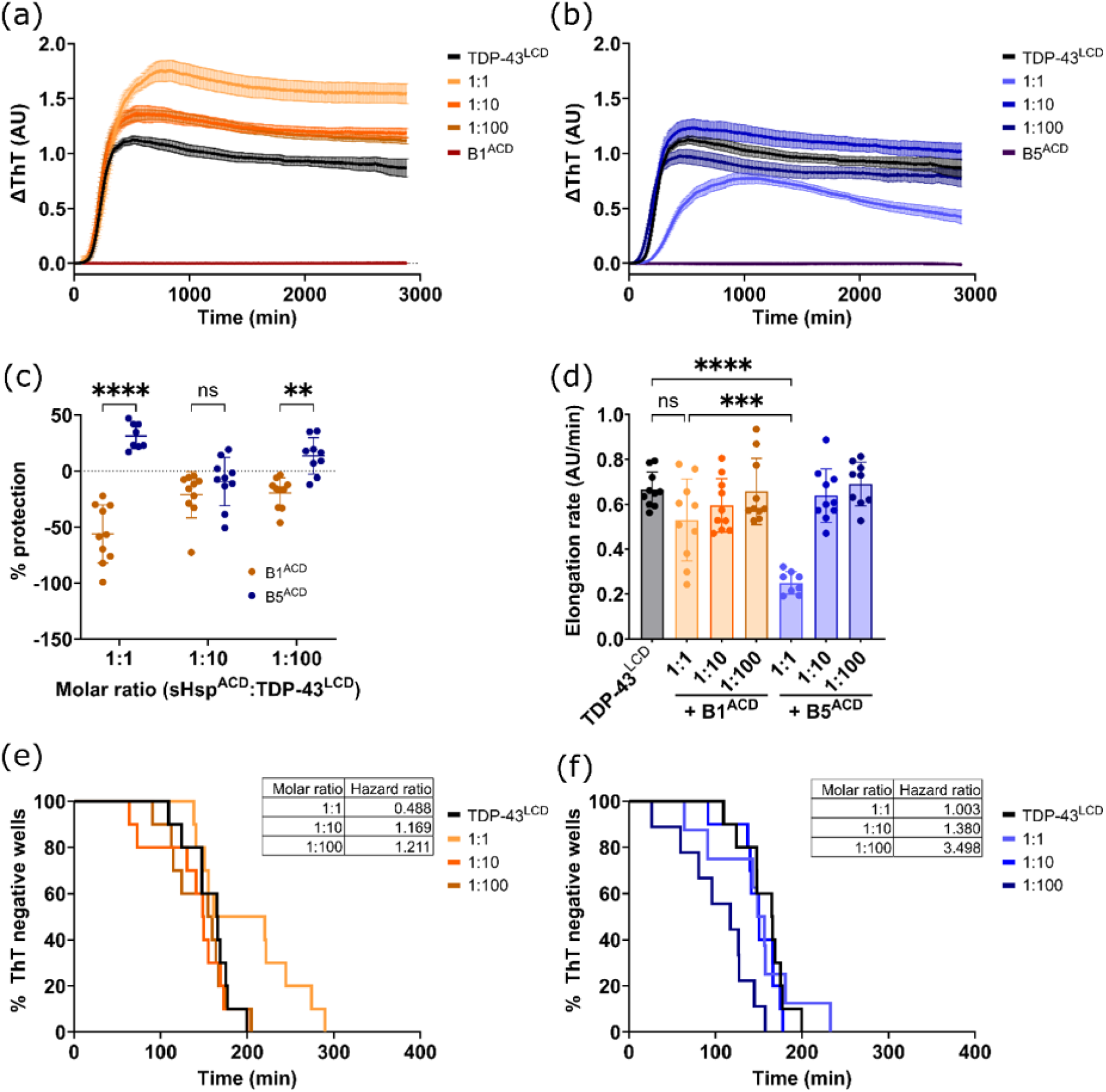
HspB1 and HspB5 partially inhibit TDP-43^LCD^ fibrillation through their α-crystallin domains. TDP-43^LCD^ (10 μM) was incubated with different molar ratios (1:1, 1:10, 1:100) of HspB1^ACD^ or HspB5^ACD^ (sHsp:TDP-43^LCD^) and 25 μM ThT at 37°C and monitored for changes in ThT fluorescence at 490 nm. Samples were shaken at 200 rpm for 30 s prior to each cycle. (a) ThT curve for TDP-43^LCD^ in the presence of HspB1^ACD^. (b) ThT curve for TDP-43^LCD^ in the presence of HspB5^ACD^. (c) Percentage protection against TDP-43^LCD^ fibrillation for HspB1^ACD^ and HspB5^ACD^. Protection was calculated relative to the mean ThT fluorescence intensity of TDP-43^LCD^ alone. (d) Elongation rate of TDP-43^LCD^ fibrils in the presence of HspB1^ACD^ or HspB5^ACD^. (e) Kaplan-Meier analysis of TDP-43^LCD^ fibrillation in the presence of HspB1^ACD^. Hazard ratio for each molar ratio is shown inset. (f) Kaplan-Meier analysis of TDP-43^LCD^ fibrillation in the presence of HspB5^ACD^. Hazard ratio for each molar ratio is shown inset. Data plotted as mean ± SEM (a, b) or mean ± SD (c, d) (n = 10 technical replicates from 2 independent experiments. Data analysed by (c) two-way ANOVA with Šidak’s post-hoc test or (d) one-way ANOVA with Tukey’s post-hoc test (** = P < 0.01, *** = P < 0.001, **** = P < 0.0001).

### HspB1 and HspB5 specifically interact with small oligomers of TDP-43^LCD^

Since both HspB1 and HspB5 delayed the formation of ThT-positive TDP-43^LCD^ aggregates, we hypothesised that they do so by interacting with monomers or small oligomers of TDP-43^LCD^. To investigate this, we performed two-colour coincidence detection (TCCD) experiments using Cy3-labelled TDP-43^LCD^ and Cy5-labelled sHsp isoforms (37); to do this, TDP-43^LCD^ was either incubated alone or in the presence of sHsp variants under identical conditions as described in the assays above. When measuring diffusion of Cy3-labelled TDP-43^LCD^ and Cy5-labelled chaperones through the confocal volume, we frequently observed large peaks in fluorescence intensity that were significantly greater than background intensity (**Fig. S2**). We hypothesised that smaller peaks in Cy3 fluorescence represent monomeric TDP-43^LCD^, while larger peaks in fluorescence intensity represent higher order states such as oligomers and larger aggregates. Notably, peaks in fluorescence intensity corresponding to Cy5-labelled chaperones often were observed at concurrently to peaks in Cy3 fluorescence (**Fig. S2**) Therefore, we sought to quantify the coincidence of fluorescence intensity peaks from Cy3-labelled TDP-43^LCD^ with those observed from Cy5-labelled chaperones, and vice versa.

When TDP-43^LCD^ was incubated with the sHsps, TDP-43^LCD^ peaks were found to be coincident with peaks from the molecular chaperones significantly more often compared to when incubated with the non-chaperone control protein, GFP, indicating that TDP-43^LCD^ interacts specifically with the chaperones (**Fig. 5a**). Conversely, sHsps peaks, but not GFP, were coincident with TDP-43^LCD^ peaks (**Fig. 5b**). Critically, the addition of 6 M guanidine-HCl to these samples resulted in a significant reduction in the size and number of TDP-43^LCD^ peaks, such that they were virtually absent, confirming that these peaks indeed represented aggregated species (**Fig. 5c, d**). TDP-43^LCD^ peak height (which is an indicator of oligomeric size) was reduced in the presence of all sHsp isoforms except HspB5^WT^, suggesting that incubation of TDP-43^LCD^ with these chaperones reduces the size of TDP-43^LCD^ oligomers (**Fig. 5c**). Furthermore, the number of TDP-43^LCD^ peaks was significantly reduced in the presence of each of the chaperones, as well as GFP, suggesting that less TDP-43^LCD^ oligomers are formed upon co-incubation (**Fig. 5d**).

**Figure 5.**
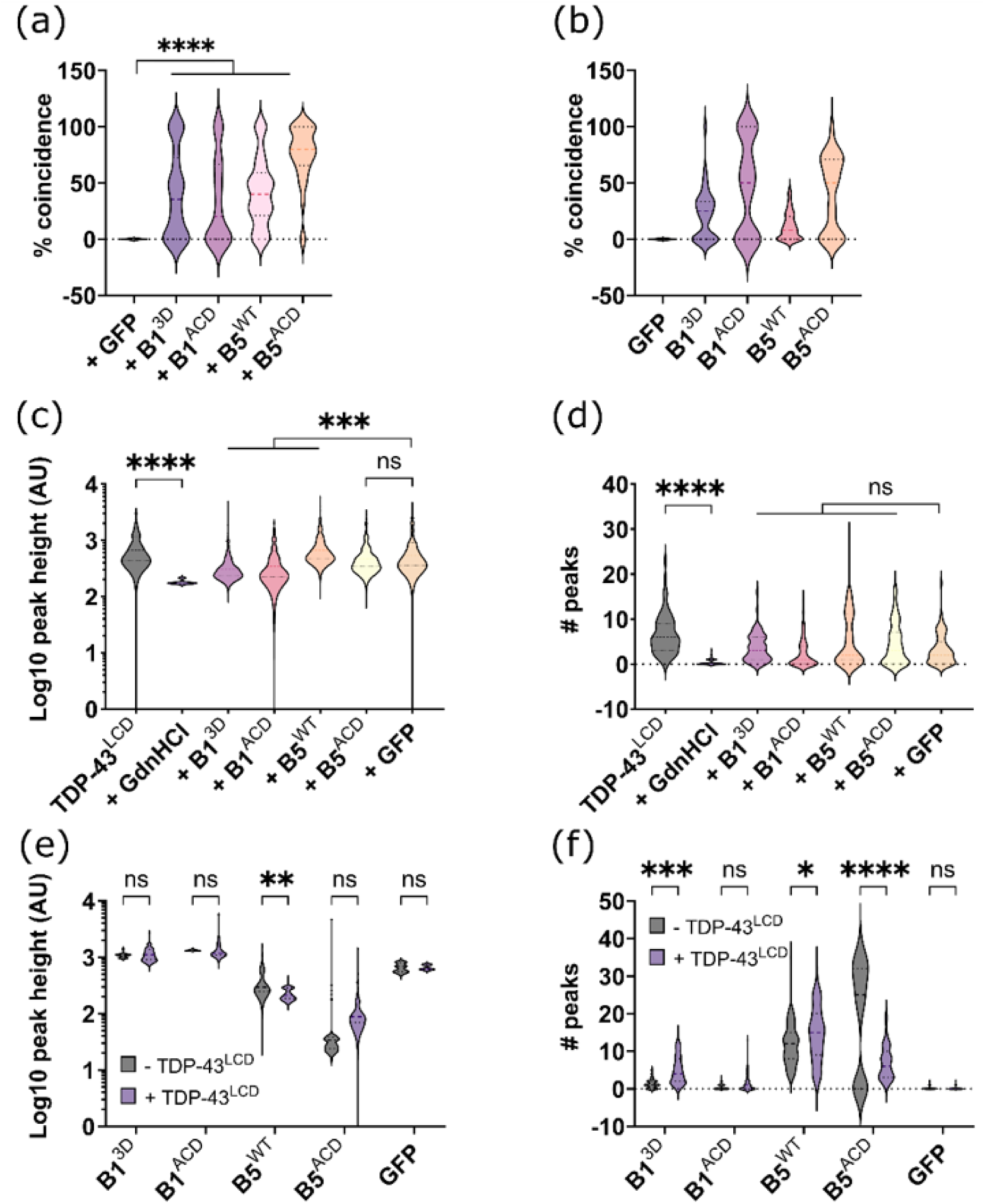
HspB1 and HspB5 specifically interact with TDP-43^LCD^ oligomers. Two-colour coincidence detection (TCCD) was performed using a Leica TCS SP8 FALCON laser scanning confocal microscope. (a) Example trace of Cy3-TDP-43^LCD^ and Cy5-HspB1^3D^. TDP-43^LCD^ is indicated in light grey, while HspB1^3D^ is indicated in dark grey. Coincident peaks are indicated by orange circles, while non-coincident peaks are indicated by blue circles. (a) Percentage coincidence of chaperones with TDP-43^LCD^. (b) Percentage coincidence of TDP-43^LCD^ with chaperones. (c) TDP-43^LCD^ peak height in the absence or presence of chaperones. (d) Number of TDP-43^LCD^ peaks in the absence or presence of chaperones. (e) Chaperone peak height in the absence or presence of TDP-43^LCD^. (f) Number of chaperone peaks in the absence or presence of TDP-43^LCD^. Data analysed by Kruskal-Wallis test with Dunn’s post-hoc test (a, c, d) or two-way ANOVA with Šidák’s post-hoc test (e, f) (n = 75 technical replicates from 3 independent experiments) (* = P < 0.05, ** = P < 0.01, *** = P < 0.001, **** = P < 0.0001). No statistical analysis was performed for (b) due to uneven labelling efficiencies.

When we looked at the fluorescence intensity originating from the chaperones that were coincident with TDP-43^LCD^, we found that the peak height was reduced only when TDP-43^LCD^ was co-incubated with HspB5^WT^ (**Fig 5e**). We observed a significant increase in the number of HspB1^3D^ and HspB5^WT^ peaks when they were in the presence of TDP-43^LCD^, suggesting that they were being recruited to TDP-43^LCD^ aggregates (**Fig. 5f**). In contrast, there were fewer HspB5^ACD^ peaks in the presence of TDP-43^LCD^ (**Fig. 5f**). Together, these results indicate that HspB1 and HspB5 specifically bind to TDP-43^LCD^ oligomers through interactions that involve their core ACDs and, by doing so, inhibit the assembly of TDP-43^LCD^ oligomers into higher order aggregates.

### Sodium chloride induces the condensation of the TDP-43^LCD^ in a concentration-dependent manner

In addition to its role in the pathological aggregation of TDP-43 in ALS and FTLD, TDP-43^LCD^ has been shown to be sufficient and necessary for TDP-43 condensation (33). Condensation of TDP-43^LCD^ is promoted by multiple factors including increased salt concentration, decreased temperature, pH, molecular crowding, and the presence of RNA (33, 38–40). We sought to characterise the salt-induced phase separation behaviour of TDP-43^LCD^ in our system. Purified TDP-43^LCD^ (20 µM) was incubated in the presence of increasing concentrations of NaCl (0-500 mM) for 1 h at room temperature before an aliquot was loaded onto a glass slide for confocal imaging. In the absence of NaCl, small and sparsely abundant droplets were observed on the glass surface (**Fig. 6a**); these were observed to become much larger and more abundant with increasing concentrations of NaCl. These droplets exhibited liquid-like behaviour - they were found to fuse with one another and wet the surface of the glass coverslip. Quantification of the protein concentration in the dilute phase (i.e. non-condensate fraction) of the condensation reactions revealed a decrease in the concentration of TDP-43^LCD^ with increasing NaCl concentrations, indicating increased incorporation of TDP-43^LCD^ into condensates (**Fig. 6b**). A kinetic light scattering assay was performed in which increasing optical density at 340 nm, 400 nm and 600 nm was used as an indication of the formation of condensates of increasing size. As expected, increasing salt concentration resulted in more rapid formation of condensates of various sizes, as the rate of condensate formation appeared lower for larger condensates (**Fig. 6c-e**).

**Figure 6.**
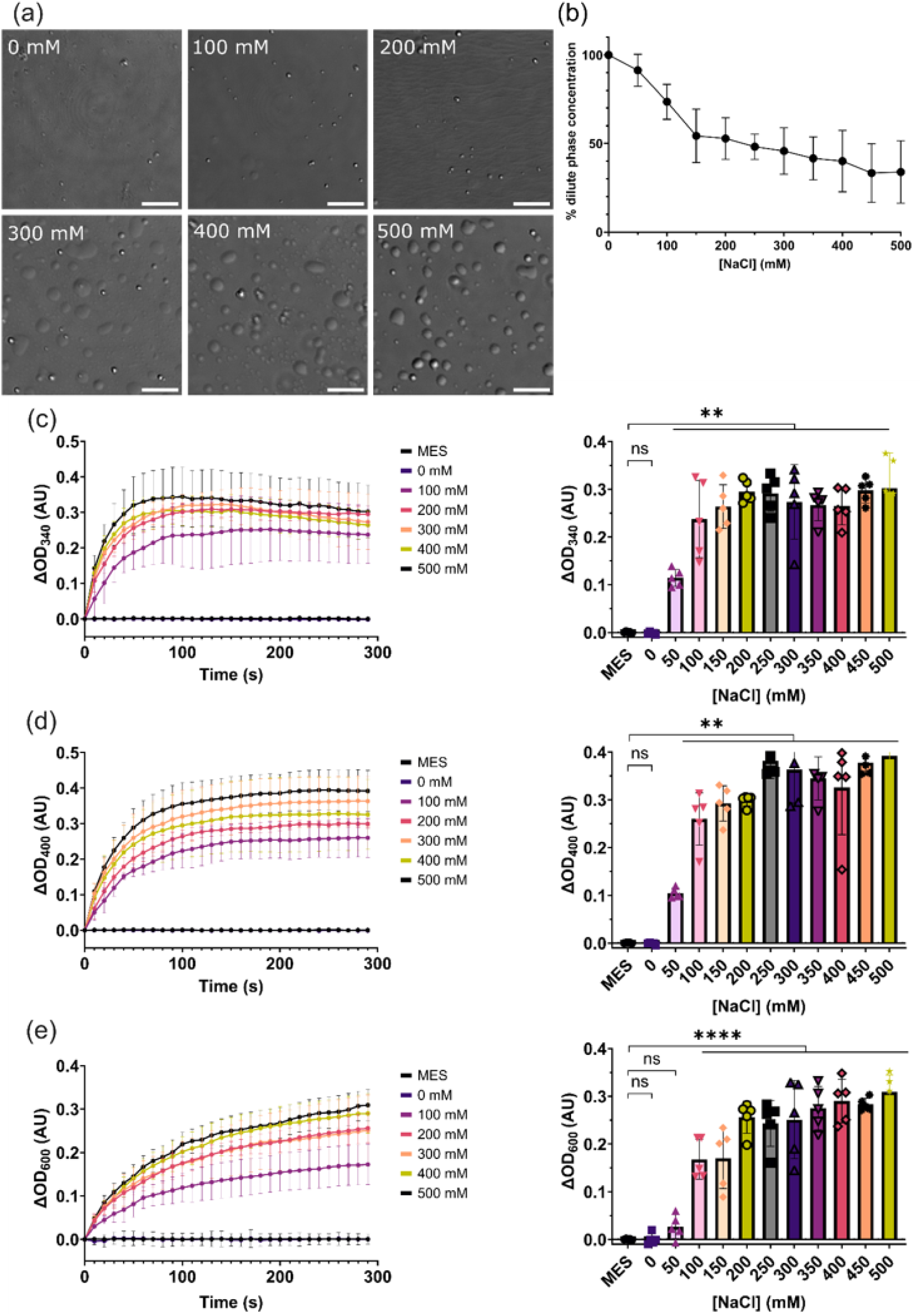
TDP-43^LCD^ phase separates in a NaCl concentration-dependent manner. TDP-43^LCD^ (20 μM) in 20 mM MES (pH 6.0) was incubated with different concentrations of NaCl (0 – 500 mM) to induce phase separation. (a) Differential interference contrast (DIC) microscopy images of TDP-43^LCD^ condensates. NaCl concentrations are indicated. Scale bars represent 10 μm. (b) Quantification of proportion of TDP-43^LCD^ that remains in the dilute phase (supernatant) following incubation. (c-e) Quantification of the change in OD at (c) 340 nm, (d) 400 nm, and (e) 600 nm following mixing of TDP-43^LCD^ with NaCl. Left panel shows change in OD as a function of time with increasing NaCl concentration. Right panel shows endpoint OD values with increasing NaCl concentration. Data plotted as mean ± SD (n = 5 technical replicates from 2 independent experiments). Data analysed by one-way ANOVA with Dunnett’s post-hoc test with comparison to MES control (** = P < 0.01, **** = P < 0.0001).

### HspB1^3D^ promotes TDP-43^LCD^ condensation, but HspB5 does not

As the sHsp HspB1 regulates the cytoplasmic condensation of TDP-43 (29), we investigated how HspB1 modifies TDP-43 phase separation *in vitro* and determined whether its homolog HspB5 also influences TDP-43 condensation. In these experiments, we used the phosphomimetic variant of HspB1, HspB1^3D^, as it was most effective at inhibiting the aggregation of TDP-43^LCD^ (see **Fig. 3**). In contrast, the wild-type form of HspB5 (HspB5^WT^) was used in these experiments, as similar phosphomimicking mutations have a less pronounced effect upon its oligomeric distribution (21, 35, 40). Surprisingly, simply mixing HspB1^3D^ with TDP-43^LCD^ at equimolar concentrations resulted in the formation of condensates, even in the absence of NaCl (**Fig. 7a**), indicating that HspB1^3D^ promotes TDP-43^LCD^ phase separation under these conditions. These condensates appeared larger and more numerous than those formed in the presence of NaCl. In contrast to the ability of HspB1^3D^ to potentiate TDP-43^LCD^ condensation, few condensates were observed when TDP-43^LCD^ was incubated with HspB5^WT^, indicating that HspB5 does not promote condensation of TDP-43^LCD^ under these conditions. Quantification of these differences via a kinetic light scattering assay revealed significantly greater turbidity at all wavelengths measured when TDP-43^LCD^ was incubated with HspB1^3D^ compared to when NaCl and/or HspB5^WT^ were present (**Fig. 7b-d**). These findings suggest that, despite their sequence and structural similarities, HspB1^3D^ and HspB5^WT^ interact differently with TDP-43^LCD^ under conditions that promote its phase separation.

**Figure 7.**
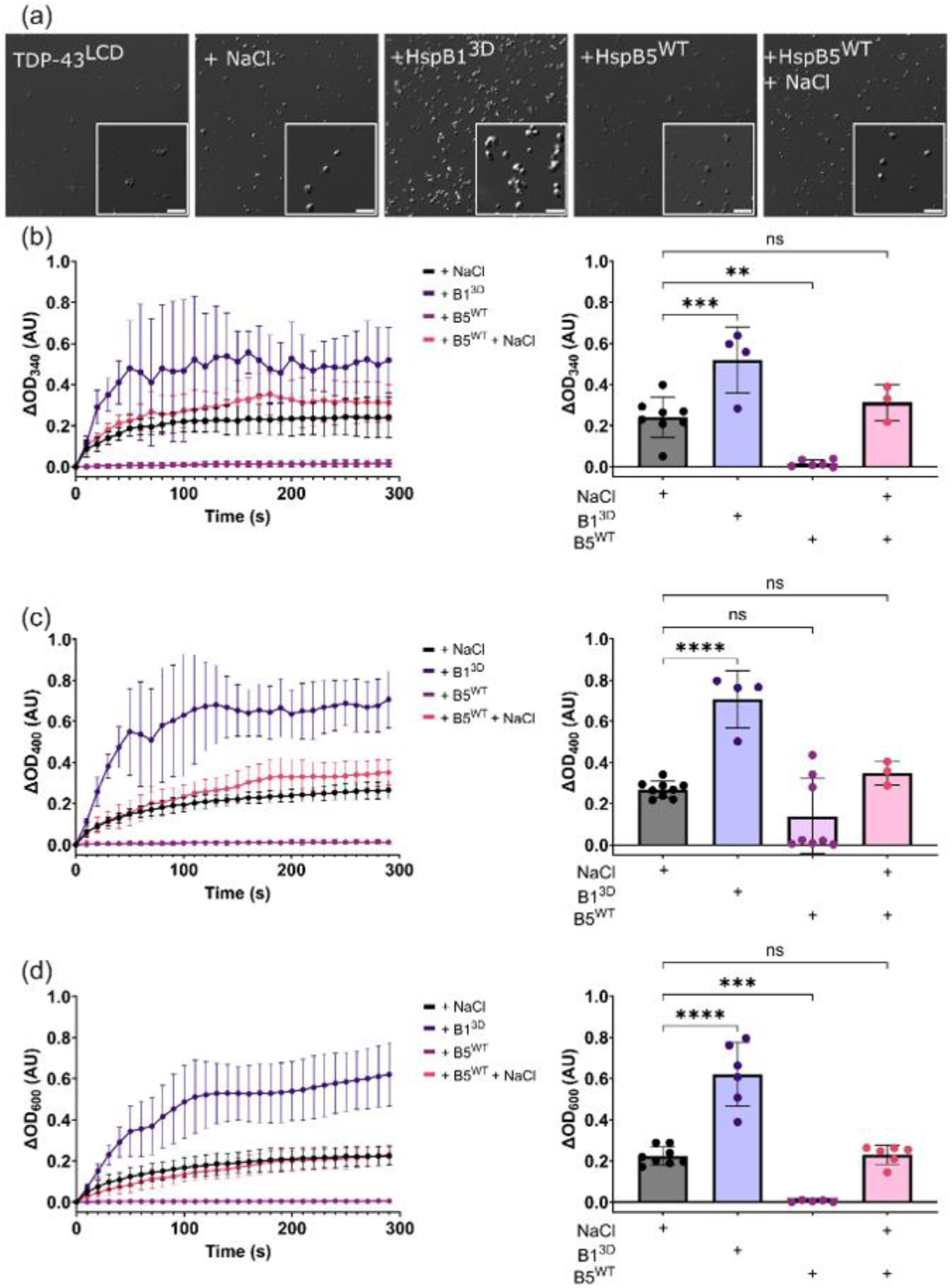
HspB1 promotes TDP-43^LCD^ phase separation. TDP-43^LCD^ (20 μM) in 20 mM MES (pH 6.0) was incubated in the presence of HspB1^3D^ (20 μM), or HspB5^WT^ (20 μM) in the presence or absence of NaCl (200 mM) (a) Differential interference contrast (DIC) confocal microscopy images of TDP-43^LCD^ condensates formed in the presence of either chaperone. Scale bars represent 5 μm. (b-d) Quantification of the change in OD at (c) 340 nm, (d) 400 nm, and (e) 600 nm following mixing of TDP-43^LCD^ with either chaperone. Left panel shows change in OD as a function of time. Right panel shows endpoint OD values. Data plotted as mean ± SD (n ≥ 3 technical replicates from 2 independent experiments). Data analysed by one-way ANOVA with Dunnett’s post-hoc test with comparison to NaCl control (** = P < 0.01, *** = P < 0.001, **** = P < 0.0001).

### The ACDs of HspB1 and HspB5 do not significantly impact TDP-43^LCD^ condensation

We sought to ascertain whether the central α-crystallin domain of HspB1 played a role in promoting the condensation of TDP-43^LCD^. To do so, TDP-43^LCD^ was incubated with HspB1^ACD^ or HspB5^ACD^, either in the absence or presence of NaCl (**Fig. 8**). Incubation of TDP-43^LCD^ with either HspB1^ACD^ or HspB5^ACD^ in the absence of NaCl resulted in the formation of small clusters of condensates that appeared more numerous than those formed by TDP-43^LCD^ alone (**Fig. 8a**). Despite this, both samples exhibited low endpoint optical density compared to TDP-43^LCD^ incubated with NaCl (**Fig. 8b-d**). In the presence of NaCl, HspB1^ACD^ and HspB5^ACD^ did not significantly alter the appearance of TDP-43^LCD^ condensate (**Fig. 8a**), nor did they alter the endpoint optical density of these samples (**Fig. 8b-d**). Thus, neither HspB1^ACD^ nor HspB5^ACD^ significantly altered the condensation of TDP-43^LCD^, consistent with them having a high degree of secondary structure that is generally considered incompatible with condensation. From these findings, we conclude that HspB1 promotes TDP-43^LCD^ phase separation via interactions involving its disordered N- and/or C-terminal regions.

**Figure 8.**
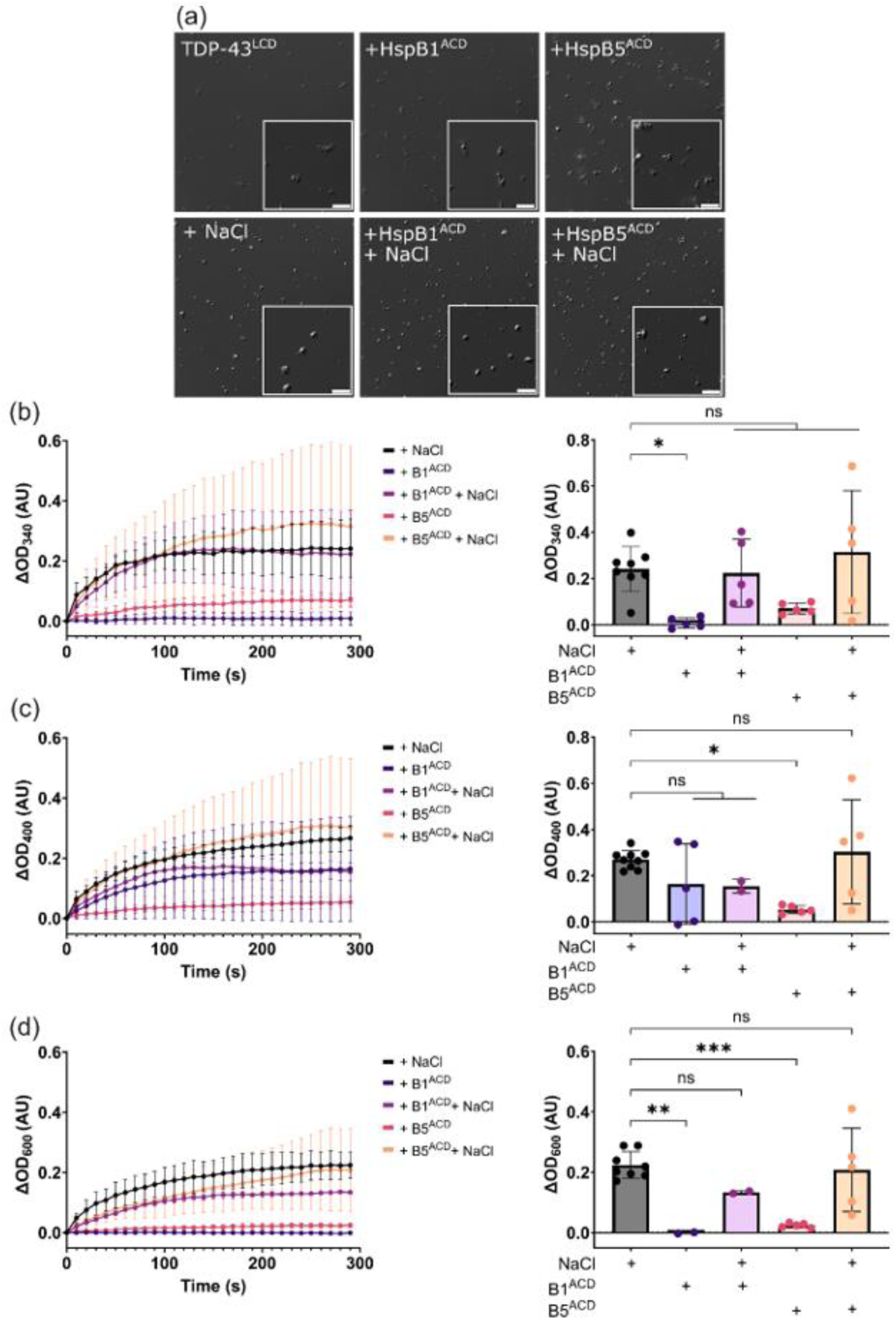
HspB1^ACD^ and HspB5^ACD^ do not significantly impact condensation of TDP-43^LCD^. TDP-43^LCD^ (20 μM) in 20 mM MES (pH 6.0) was incubated with HspB1^ACD^ (20 μM) or HspB5^ACD^ (20 μM) in the presence or absence of NaCl (200 mM). (a) Differential interference contrast (DIC) confocal microscopy images of TDP-43^LCD^ condensates formed in the presence of either chaperone. Scale bars represent 5 μm. (b-d) Quantification of the change in OD at (c) 340 nm, (d) 400 nm, and (e) 600 nm following mixing of TDP-43^LCD^ with either chaperone. Left panel shows change in OD as a function of time. Right panel shows endpoint OD values. Data plotted as mean ± SD (n ≥ 2 technical replicates from 2 independent experiments). Data analysed by one-way ANOVA with Dunnett’s post-hoc test in comparison to NaCl control (* = P < 0.05, ** = P < 0.01, *** = P < 0.001).

### HspB1 and HspB5 partition into TDP-43^LCD^ condensates

HspB1 has been reported to partition within cytoplasmic TDP-43 condensates, where it interacts with the RNA binding domain and LCD of TDP-43 (30). Therefore, we investigated whether the effects of HspB1 and HspB5 upon TDP-43^LCD^ condensation correlates with their incorporation into condensates. To this end, we utilised Cy3-labelled TDP-43^LCD^ and Cy5-labelled full-length or ACD isoforms of the sHsps to investigate their possible localisation in these condensates. In doing so, we calculated the free energy of partitioning (ΔG_partition_) of samples as a measure of the proportion of each protein within condensates, as described previously (41). We observed partitioning of HspB1^3D^, HspB1^ACD^, HspB5^WT^ and HspB5^ACD^ within TDP-43^LCD^ condensates (**Fig. 9**). In contrast, mCherry, a non-chaperone control protein, was excluded and remained outside condensates (**Fig. S3**). Partitioning of TDP-43^LCD^ to condensates did not vary significantly over time, indicating that it rapidly reaches a steady state when induced to phase separate (**Fig. 9a**). Interestingly, there was a time-dependent increase in the partitioning of HspB1^3D^ within condensates, indicating that the interaction between TDP-43^LCD^ and HspB1^3D^ is enhanced over time (**Fig. 9b**). All other isoforms tested initially partitioned into TDP-43^LCD^ condensates, with both ACD forms doing so the most, and then decreased over time (**Fig. 9c-e**). These findings indicate that the partitioning of HspB1 and HspB5 is specific and dynamic.

**Figure 9.**
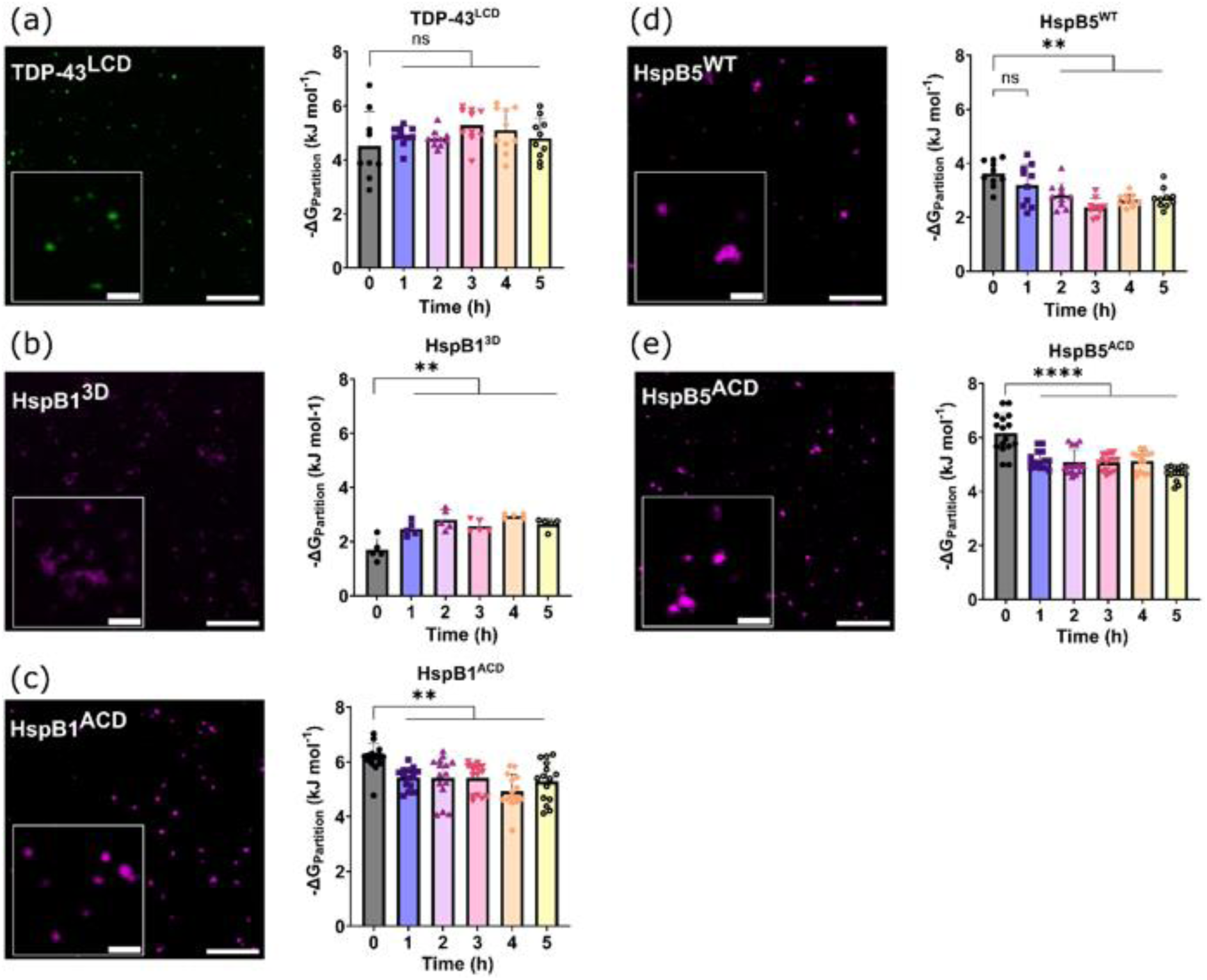
HspB1 and HspB5 partition specifically within TDP-43^LCD^ condensates. Cy3-labelled TDP-43^LCD^ or unlabelled TDP-43^LCD^ (20 μM) was induced to phase separate in the presence of Cy5-labelled forms of HspB1^3D^, HspB1^ACD^, HspB5^WT^ or HspB5^ACD^. Images were taken at 1 h intervals at 40× magnification using a Leica TCS SP5 laser scanning confocal microscope. Microscopy images were used to calculate the partitioning of each protein to TDP-43^LCD^ condensates. (a-e) Partitioning of (a) Cy3-TDP-43^LCD^, (b) Cy5-HspB1^3D^, (c) Cy5-HspB1^ACD^, (d) Cy5-HspB5^WT^, (e) Cy5-HspB5^ACD^ to condensates. Left panels show confocal microscopy images of protein partitioning to TDP-43^LCD^ condensates at t = 0. Scale bars represent 10 μm. Zoomed images are shown inset. Scale bars for inset images represent 2 μm. Right panels show -ΔGPartition of fluorescently labelled proteins to condensates over time. Data plotted as mean ± SD (n ≥ 5 technical replicates from at least 2 independent experiments). Data analysed by one-way ANOVA with Dunnett’s post-hoc analysis with comparison to t = 0 (** = P < 0.01, **** = P < 0.0001).

### Both HspB1^3D^ and HspB5^WT^ maintain the dynamic nature of TDP-43^LCD^ condensates

We next investigated how HspB1^3D^ and HspB5^WT^ alter the dynamic properties of TDP-43^LCD^ condensates. To this end, we performed fluorescence recovery after photobleaching (FRAP) (42). In these experiments, the entire area of a condensate was bleached to evaluate the movement of TDP-43^LCD^ protein into condensates. By plotting the fluorescence recovery over time, we derived two key parameters: (i) the mobile fraction, which is the fraction of TDP-43^LCD^ capable of exchange and (ii) half-time, which is the time taken to reach half-maximal fluorescence recovery and is an indicator of the speed at which TDP-43^LCD^ undergoes exchange. Here, we observed distinct differences in the dynamic exchange of TDP-43^LCD^ within condensates and between the condensed and dilute phases when in the presence or absence of HspB1^3D^ and HspB5^WT^ (**Fig. 10**). In the absence of either sHsp, fully bleached TDP-43^LCD^ condensates exhibited poor fluorescence recovery, yielding a low mobile fraction and half-time to recovery; notably, this continued to decline for condensates at later stages of incubation (**Fig. 10a, c, d**). In the presence of HspB1^3D^, the mobile fraction of bleached TDP-43^LCD^ condensates increased, indicating that a greater proportion of TDP-43^LCD^ within condensates were capable of exchanging with the surrounding solution (**Fig. 10c**). Although the mobile fraction of these condensates declined at later stages of incubation, they remained significantly more mobile than those containing only TDP-43^LCD^. Further, the half-time to recovery for these condensates was reduced in the presence of HspB1^3D^, indicating faster exchange of TDP-43^LCD^ between the two phases (**Fig. 10d**). This effect was maintained across the time course of the assay. In contrast, HspB5^WT^ did not alter the proportion of TDP-43^LCD^ capable of exchange between condensates and the surrounding solution; rather it inhibited the loss of dynamic exchange over time, such that after 5 h condensates that contained both TDP-43^LCD^ and HspB5^WT^ had a significantly greater mobile fraction than those composed of TDP-43^LCD^ alone (**Fig. 10c**). These condensates also maintained a similar half-time to recovery over the 5 h period, such that it was greater than that of condensates made of TDP-43^LCD^ alone after 5 h (**Fig. 10d**). These data suggest that the presence of HspB5^WT^ inhibits reductions in TDP-43^LCD^ mobility between the condensed and dispersed phases.

**Figure 10.**
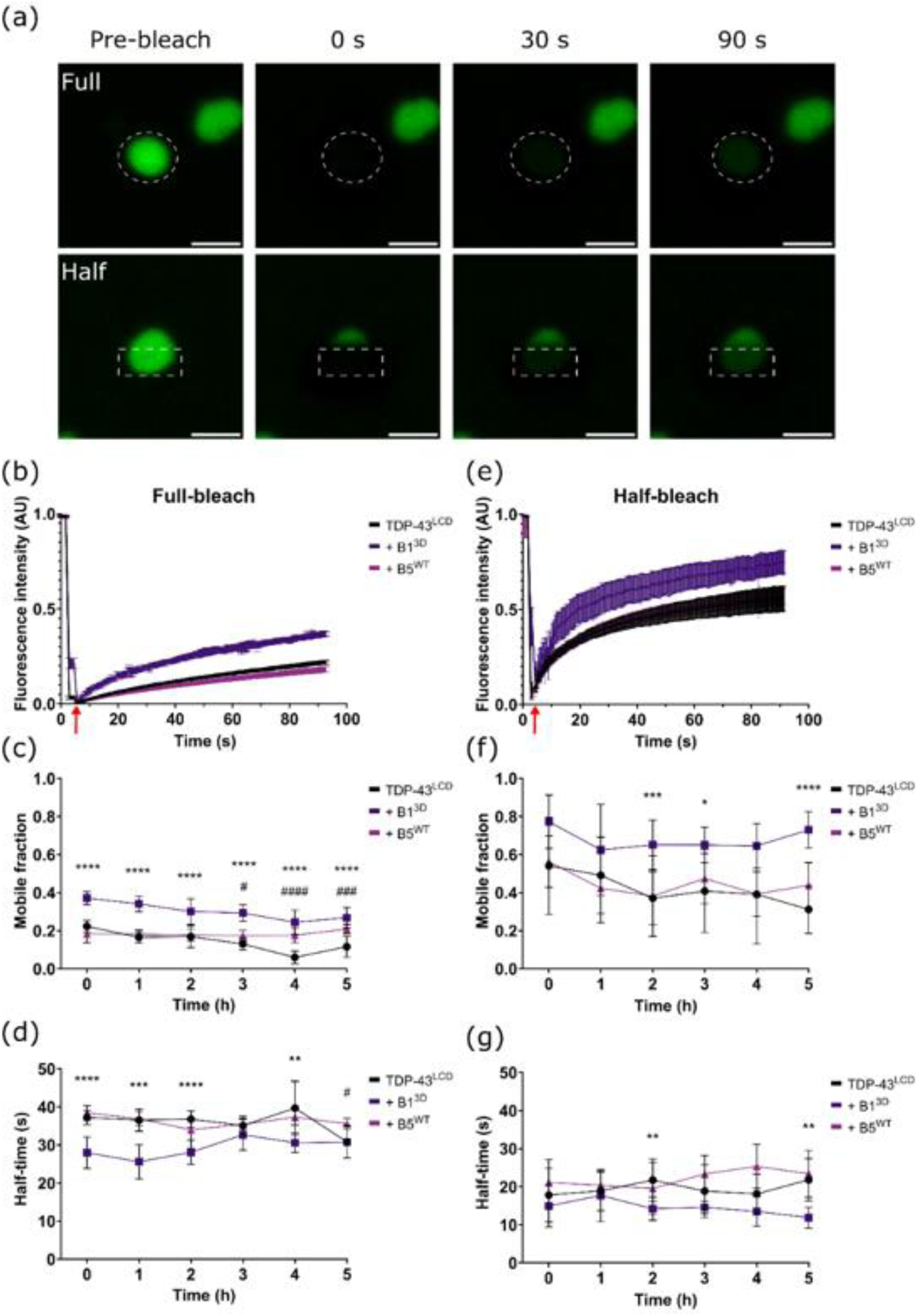
HspB1 and HspB5 differentially alter the dynamic properties of TDP-43^LCD^ condensates. Cy3-labelled TDP-43^LCD^ was induced to phase separate in the presence of Cy3-labelled forms of HspB1^3D^ or HspB5^WT^ and evaluated by FRAP. The fluorescence recovery of fully bleached or half-bleached TDP-43^LCD^ condensates was monitored over ∼90 s and analysed using EasyFRAP to calculate the mobile fraction and half-time to recovery. (a) Example images showing fluorescence recovery of photobleached TDP-43^LCD^ condensates. Top row shows a fully bleached condensate. Bottom row shows a half-bleached condensate. Scale bars represent 2 μm. (b) FRAP curve for fully bleached TDP-43^LCD^ condensates in the absence or presence of HspB1^3D^ or HspB5^WT^ at t = 0 h. Red arrow shows the time at which photobleaching was performed. (c) Mobile fraction of fully bleached TDP-43^LCD^ condensates. (d) Half-time to recovery of fully bleached TDP-43^LCD^ condensates. (e) FRAP curve for half-bleached TDP-43^LCD^ condensates in the absence or presence of HspB1^3D^ or HspB5^WT^ at t = 0 h. Red arrow shows the time at which photobleaching was performed. (f) Mobile fraction of half-bleached TDP-43^LCD^ condensates. (g) Half-time to recovery of half-bleach TDP-43^LCD^ condensates. Data plotted as mean ± SEM (b, e) or mean ± SD (c, d, f, g) (n ≥ 5 technical replicates from three independent experiments). Data analysed by two-way ANOVA with Dunnett’s post-hoc test with comparison to TDP-43^LCD^ control. (*/# = P < 0.05, ** = P < 0.01, ***/### = P < 0.001, ****/#### = P <0.0001. Asterisks indicate significant differences for condensates in the presence of HspB1^3D^ compared to TDP-43^LCD^ alone. Hash symbols indicate significant differences for condensates in the presence of HspB5^WT^ compared to TDP-43^LCD^ alone).

We also evaluated the dynamic exchange of protein within the condensed phase via FRAP. To do so, approximately half of a condensate was bleached to evaluate the movement of TDP-43^LCD^ protein within individual condensates (**Fig. 10e**). For condensates containing TDP-43^LCD^ alone, we observed a decrease in the mobile fraction across the course of 5 h, indicative of a reduction in the internal dynamics of these condensates over time (**Fig. 10f**). There was no significant change in their half-time to recovery over time (**Fig. 10g**). In the presence of HspB1^3D^, the mobile fraction of TDP-43^LCD^ was enhanced compared to that of condensates containing TDP-43^LCD^ alone (**Fig. 10f**). The enhancement of TDP-43^LCD^ mobility within condensates by HspB1^3D^ was maintained after 5 h. Although the half-time to recovery was not initially altered by the presence of HspB1^3D^, after 5 h the half-time to recovery was significantly lower than that of condensates containing only TDP-43^LCD^, indicating that the presence of HspB1^3D^ enhanced the rate of TDP-43^LCD^ mobility throughout condensates (**Fig. 10g**). HspB5^WT^ did not significantly alter the mobile fraction at any point, indicating that its presence does not alter the proportion of TDP-43^LCD^ capable of exchange within condensates (**Fig. 10f**). However, it did cause a significant increase in the half-time to recovery of condensates at 4 h, indicative of more rapid exchange (**Fig. 10g**). Collectively, these findings indicate that HspB1 and HspB5 differentially alter the dynamic exchange of TDP-43^LCD^ between the condensed and dispersed phases and within condensates themselves, thereby contributing to the dynamics of TDP-43^LCD^ condensates.

## Discussion

While condensates and amyloids can arise via distinct protein interactions, aggregation via condensation is primarily driven by protein regions that can engage in multimodal interactions (43). Invariably, these regions are intrinsically disordered and, thus, sample conformationally heterogeneous states. Therefore, the cellular mechanisms that maintain the dynamic state of protein condensates will differ from those that maintain proteins in a soluble state. The capacity of sHsps to inhibit the assembly of amyloid-like fibrils is well-established (10, 16, 44), however much less is known about how sHsps engage with potential client proteins during condensation. In this study, we show that HspB1 and HspB5 differentially interact with TDP-43^LCD^ during its condensation and aggregation. Notably, we demonstrate that HspB1 promotes the spontaneous condensation of TDP-43^LCD^, further emphasising the distinct interactions between different sHsps and the same client protein.

The functional differences between HspB1 and HspB5 may be explained by sequence (and structural) differences that exist throughout their amino acid sequences, particularly in the variable N- and C-terminal domains. Although full-length isoforms of HspB1 and HspB5 exhibit the generic ability to inhibit the fibrillar aggregation of TDP-43^LCD^, similar examination of their central ACDs revealed opposite effects upon TDP-43^LCD^ aggregation. Furthermore, whilst full length HspB1 promoted TDP-43^LCD^ condensation and enhanced the dynamic properties of TDP-43^LCD^ condensates, but the isolated ACD did not significantly impact TDP-43^LCD^ condensation despite it (and the ACD of HspB5) partitioning into condensates. Despite being comparatively less potent than their full-length isoforms, the isolated ACDs of HspB1 and HspB5 were both found to specifically interact with small oligomers of TDP-43^LCD^, indicating a role for this region in client binding. These findings indicate that the divergences in the functional interactions of HspB1 and HspB5 with TDP-43^LCD^ are primarily derived from differences in the N- and/or C-terminal regions, which share 38% and 37% sequence similarity, respectively. Indeed, this is consistent with previous work demonstrating that binding of various clients by HspB1 and HspB5 involves distinct regions primarily situated in the variable N- and/or C-terminal domains that flank the central α-crystallin domain (26–29, 45–47). Together, these findings suggest that HspB1 and HspB5 engage with TDP-43 via multivalent interactions.

Although this study did not examine which regions in TDP-43^LCD^ are bound by HspB1 and HspB5 during condensation and fibrillation, previous work has revealed structural features within TDP-43^LCD^ that govern these behaviours. Condensation of TDP-43^LCD^ is driven primarily by aromatic amino acids flanked by glycine and serine residues, which contribute to condensation through intermolecular π-stacking and cation-π interactions (38). Interestingly, similar aromatic motifs are present in the disordered N-terminal region of HspB1, suggesting that TDP-43^LCD^ condensation promoted by HspB1 may occur via intermolecular interactions involving these motifs on both proteins. Furthermore, TDP-43^LCD^ harbours a transient α-helical region that is crucial for condensation and undergoes structural transformation to a β-sheet during fibrillation, forming the fibril core (32, 48). Given their potent ability to recognise and bind misfolded proteins, it is likely that HspB1 and HspB5 bind this amyloidogenic core region within TDP-43^LCD^ to prevent its assembly into fibrils. Further investigation of the molecular mechanisms of these interactions may guide the development of therapeutics against TDP-43 proteinopathies.

The oligomeric state of sHsps influences their chaperone activity (21, 23–25) and this is altered by stress-induced phosphorylation of serine residues that reside within the N-terminal domain of both HspB1 and HspB5 (19–24). Mutations that mimic phosphorylation events (i.e. in the case of HspB1, at serine residues 15, 78 and 82) cause HspB1 to dissociate from large oligomers to dimers (24). Similar mutations in HspB5 (at serine residues 19, 45 and 59) have a far less pronounced effect upon its oligomeric distribution (21, 35, 41). The increase in chaperone activity because of mutations that mimic phosphorylation was strongly reflected in our findings, as the phosphomimetic mutant, HspB1^3D^, was more effective at inhibiting TDP-43^LCD^ aggregation than the wild-type isoform. Notably, the cellular conditions that promote sHsp phosphorylation are the same conditions that promote TDP-43 condensation and aggregation (30), indicating that the consequent dissociation of HspB1 into dimers functions to enhance its affinity for misfolded TDP-43. Furthermore, the enhanced dynamics of TDP-43^LCD^ in the presence of phosphomimetic HspB1^3D^ may arise from the increased negative charge of the sHsp, which promotes greater mobility of TDP-43^LCD^ through charge repulsion. Under cellular conditions, these interactions may in fact promote TDP-43 solubility, thus providing explanation for its role in the disassembly of TDP-43 condensates upon resolution of stress (30, 50).

Insights from patient-derived tissue also reveal the therapeutic potential of HspB1 and HspB5. Proteomics analysis of sporadic ALS brain and spinal cord tissue has revealed that the levels of HspB1 and HspB5 are negatively correlated with the severity of TDP-43 pathology (50). Furthermore, transcriptomic analysis of iPSC-derived motor neurons from sporadic ALS patients has shown that HspB1, but not HspB5, is upregulated during periods of oxidative stress in which TDP-43 assembles into cytoplasmic condensates (30). In contrast, the absence of HspB5 in these analyses suggests that HspB5 may fail to be transcriptionally upregulated by the onset of disease pathology in ALS motor neurons, which is reflective of the broader deficiency of proteostasis that is characteristic of ALS (51). The promising results presented in this study demonstrate the potent ability of HspB5 to delay TDP-43^LCD^ aggregation and maintain TDP-43^LCD^ condensate dynamics, which suggests that HspB5, along with HspB1, may be a viable therapeutic target for delaying ALS disease progression. Indeed, overexpression of HspB1 in the motor neurons of transgenic SOD1 G93A mice was found to inhibit disease progression (52), demonstrating its potential as a broad protectant against protein aggregation in ALS.

Protein condensation and fibrillation are both potential, sometimes overlapping, pathways that may lead to the formation of protein inclusions in neurodegenerative diseases. Therefore, characterising the mechanistic differences in these processes and the cellular systems that regulate them will be instrumental in our understanding of diseases such as ALS and FTLD-TDP. Our findings that TDP-43 condensation and fibrillation are differentially regulated by chaperones from the same class suggests that other proteins implicated in neurodegenerative diseases may also be regulated in distinct manners. As such, the design of therapeutics must reflect the understanding that condensation and fibrillation are both potential pathways that can lead to the formation of protein inclusions. In conclusion, this study demonstrates the potent ability of HspB1 and HspB5 to modify the condensation and aggregation of TDP-43. The insights provided by these findings will facilitate the advancement of therapeutics for the treatment of ALS and other neurodegenerative diseases.

## Author contributions

**Thomas B. Walker:** investigation; methodology; analysis; writing – original draft; writing – review and editing. **Joshua Trowbridge:** investigation; methodology; analysis; writing – review and editing. **Shannon McMahon**: investigation; methodology; writing – review and editing. **Nicholas Marzano**: analysis; writing – review and editing. **Lauren Rice**: investigation; methodology; writing – review and editing. **Justin J. Yerbury:** supervision; funding acquisition. **Heath Ecroyd:** conceptualisation; supervision writing – review and editing. **Luke McAlary:** conceptualisation; supervision; writing – review and editing; project administration.

## Acknowledgements

This work was funded by an NHMRC Investigator Grant awarded to Justin J. Yerbury (APP1194872). Luke McAlary also acknowledges funding from Motor Neuron Disease Australia and FightMND. We acknowledge the staff of Molecular Horizons at the University of Wollongong for their technical and administrative support.

## Materials and methods

### Preparation of bacterial expression vectors

The pJ411 bacterial expression vector for the expression of the TDP-43 LCD (TDP-43^LCD^) was a kind gift from Professor N. Fawzi (Brown University, Providence, RI; Addgene #98669). The pET3a and pET24a bacterial expression vectors containing the human *HSPB1* (HspB1) and *HSPB5* (HspB5) were used for expression of recombinant wild-type proteins (24). A variant of HspB1 designed to mimic phosphorylation at serines 15, 78 and 82 was generated by site-directed mutagenesis of serine residues to aspartic acid to produce HspB1^3D^ (24). The pET28 bacterial expression vectors for the expression of the core domains of HspB1 (residues 86-169, HspB1^ACD^) and HspB5 (residues 68-162, HspB5^ACD^) were a kind gift from Professor A. Laganowsky (Texas A&M Health Science Center). Details of all bacterial expression plasmids are shown in Supplementary Table 1. All recombinant proteins were transformed and expressed in *Escherichia coli* BL21(DE3) cells and purified as previously described (29, 53) or using a modification of a previously published protocol (33). Protein molecular weight standards used in gel electrophoresis were Protein Precision Plus Dual Color obtained from Bio-Rad Laboratories. All other chemicals, including ThT, were obtained from Sigma Aldrich, unless otherwise stated. Protein concentrations were determined using a Nanodrop 2000c spectrophotometer (ThermoFisher Scientific, Waltham, MA) based upon extinction coefficient values of 19480 M^-1^ cm^-1^ for TDP-43^LCD^, 37587 M^-1^ cm^-1^ for HspB1, 8605 M^-1^ cm^-1^ for HspB1^ACD^, 16732 M^-1^ cm^-1^ for HspB5, and 1490 M^-1^ cm^-1^ for HspB5^ACD^ as calculated using the ExPASy ProtParam tool (54).

### Expression and purification of TDP-43^LCD^

The basis for the protocols describing TDP-43^LCD^ expression and purification have been published previously (33). In this study, the protocol was modified for the generation of non-isotopically labelled TDP-43^LCD^ protein from inclusion bodies. Initial 10 mL LB media starter cultures containing 50 μg/mL kanamycin were inoculated with a single colony of transformed pJ411 TDP-43_LCD BL21(DE3) *E. coli* and incubated overnight with orbital shaking at 180 rpm at 37°C. Each 10 mL overnight starter culture was used to inoculate 1 L of 2YT media containing 50 μg/mL kanamycin. Main cultures were grown at 37°C with orbital shaking at 200 rpm until an OD_600_ of 0.6 was reached, at which point isopropyl β-D-1 thiogalactopyranoside (IPTG) was added to a final concentration of 0.5 mM. After induction, cultures were incubated for 6 h at 37°C with orbital shaking at 200 rpm. Bacteria were then harvested by centrifugation at 6000×*g* for 10 min at 4°C and stored at −20°C overnight. Bacterial pellets were resuspended in 50 mM Tris-base, 500 mM NaCl, 20 mM imidazole (pH 8.0) (25 mL per litre of main culture) with the addition of one tablet of EDTA-free protease inhibitor and lysozyme and phenymethylsulfonyl fluoride (PMSF) to final concentrations of 0.5 mg/mL and 2 mM, respectively. Cells were incubated for 30 min with gentle rocking at room temperature before being lysed using a sonicator at 10% amplitude for 20 s, five times. Insoluble material containing TDP-43^LCD^ was pelleted by centrifugation at 40 000×*g* for 20 min at 4°C and resuspended in 50 mM Tris-base, 500 mM NaCl, 20 mM imidazole, 2% (v/v) Triton X-100 (pH 8.0) (10 mL per pellet). This was repeated twice more, before being resuspended in 50 mM Tris-base, 1 M NaCl, 20 mM imidazole (pH 8.0) (10 mL per pellet) and incubated for 30 min at 4°C with gentle rocking. The lysate was pelleted by centrifugation at 40 000×*g* for 20 min at 4°C. The pellet was resuspended in filtered and degassed 20 mM Tris-base, 500 mM NaCl, 20 mM imidazole, 8 M urea (pH 8.0) (25 mL per pellet) and centrifuged at 40 000×*g* for 20 min at 4°C. The supernatant was decanted and filtered using a 0.22 μm filter and either subjected to nickel-affinity chromatography or stored at −80°C for subsequent purification.

Filtered TDP-43^LCD^ samples were injected onto a HisTrap™ column equilibrated in binding buffer containing 20 mM Tris-base, 500 mM NaCl, 20 mM imidazole, 8 M urea (pH 8.0) at a flow rate of 2 mL/min. Once the sample had completely run through, the column was washed with binding buffer at a flow rate of 3 mL/min to remove unbound proteins from the column. The protein was eluted with a gradient of 20-500 mM imidazole at a flow rate of 2.5 mL/min and collected in fractions of 10 mL volume. Fractions containing pure TDP-43^LCD^ were determined by SDS-PAGE, pooled and aliquots frozen at −80°C for later use.

### Expression and purification of sHsps and their ACDs

The expression and purification of HspB1^WT^, HspB1^3D^, HspB1^ACD^, HspB5^WT^ and HspB5^ACD^ were performed as described previously (29).

### SDS-PAGE analysis of TDP-43^LCD^ purity

SDS-PAGE analysis was used to evaluate the expression and purity of TDP-43^LCD^ protein. A 10% (v/v) acrylamide resolving gel was poured into a MINI PROTEAN® system (Bio-Rad, USA) and overlayed with 70% ethanol. Resolving gels were allowed to set for 20 min, after which the overlayed 70% ethanol was removed by absorption into paper towel. A 4% (v/v) stacking gel mix was then overlayed, and a gel comb inserted to form wells. The stacking gel was allowed to set for 20 min. Protein samples were mixed with an appropriate volume of reducing SDS-PAGE sample buffer (final concentrations: 10% (v/v) glycerol, 2% SDS, 0.1% bromophenol blue, 1.25% (v/v) β-mercaptoethanol, 62.5 mM Tris-HCl pH 6.8), and boiled for 5 min, after which samples were loaded into wells. The gel was then run at 150 V in 1× SDS running buffer (192 mM glycine, 3.5 mM SDS, 25 mM Tris) for 1 h. Afterwards, the gel was microwaved in MilliQ water for 30 s at maximum power and incubated at room temperature for 5 min with gentle rocking. This was repeated once more with fresh MilliQ water, before protein bands were visualised by incubation in Coomassie Brilliant Blue G-250 stain solution (Sigma Aldrich) for 15-30 min at room temperature with gentle rocking. Gels were destained by incubating them in MilliQ water for 30 min at room temperature with gentle rocking until proteins were clearly resolved. Protein bands of interest were identified using Precision Plus Protein™ dual colour molecular weight markers (Bio-Rad, USA).

### Buffer exchange of proteins

Purified proteins were buffer exchanged into 20 mM MES (pH 6.0) prior to all analyses using Zeba Spin 7000 MWCO desalting columns (Sigma Aldrich). Briefly, spin columns were centrifuged at 1500×*g* for 1 min to remove the storage solution. Columns were equilibrated by adding 20 mM MES (pH 6.0) and centrifuging at 1500×*g* for 1 min. This was repeated 4 times. Samples were aliquoted into the spin columns and buffer exchanged into 20 mM MES (pH 6.0) via centrifugation at 1500×*g* for 2 min (for TDP-43^LCD^) or 30 s (HspB1 and HspB5) into a low-protein binding microcentrifuge tube. Samples were immediately centrifuged at 16 300×*g* for 5 min to pellet any insoluble material. The supernatant was transferred to a fresh low-protein binding microcentrifuge tube and diluted to the desired concentration.

### Thioflavin-T assay

The formation of TDP-43^LCD^ aggregates was monitored using an *in situ* thioflavin-T (ThT) binding assay. Briefly, purified TDP-43^LCD^ protein was buffer exchanged from 8M urea into 20 mM MES (pH 6.0) as described above. The protein was immediately centrifuged to pellet any particulate material and the supernatant was transferred to a fresh low-protein binding microfuge tube and its concentration determined. Varying concentrations (50, 20, 10, 5, and 2.5 μM) of TDP-43^LCD^ were incubated with 25 μM ThT in 20 mM MES (pH 6.0) at room temperature and loaded into a clear-bottomed low-protein binding 384-well plate (Greiner, Germany). The plate was incubated in a PolarStar Omega Plate Reader (BMG Labtechnologies, Australia) at 37°C for 30 min before being sealed with adhesive film. Excitation and emission filters were set at 440 and 490 nm, respectively. The plate underwent double orbital shaking at 200 rpm for 30 s at the beginning of a 350 s cycle for at least 180 cycles, totalling ∼72 h.

The formation of TDP-43^LCD^ aggregates in the presence of sHsps was also monitored using an *in situ* ThT binding assay. Briefly, 10 μM TDP-43^LCD^ was incubated with varying concentrations (50, 10, 2, 1, or 0.2 μM) of either HspB1 or HspB5 with 25 μM ThT in 20 mM MES (pH 6.0) at room temperature. The reaction mixtures were loaded into a clear-bottomed low-protein binding 384-well plate (Greiner, Germany) and incubated in a PolarStar Omega Plate Reader (BMG Lab Technologies, Australia) at 37°C for 30 min before being sealed with adhesive film. The plate underwent double orbital shaking at 200 rpm for 30 s at the beginning of a 350 s cycle for at least 742 cycles, with ThT fluorescence measured by excitation at 450 nm and emission read at 480 nm using the bottom optic of the plate reader.

The kinetics of aggregation were analysed by deriving the length of the lag-phase and fibril elongation rate from the equation for the Boltzmann sigmoidal curve. The main parameters used in this equation are final fluorescence (*F_f_*), initial fluorescence (*F_i_*), time to half maximal fluorescence (*t_50_*), and the slope of the curve (*k*). The following equations have been described previously (55).

The equation of the Boltzmann-sigmoidal curve is given as:

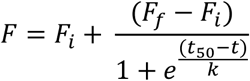

The slope of the line at any point is:

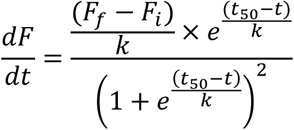

When *t* = *t_50_* (i.e. the inflexion point, when the elongation rate is maximal), the elongation rate is:

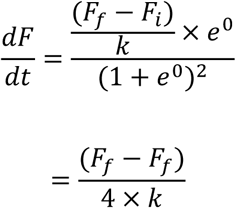

The lag phase is when:

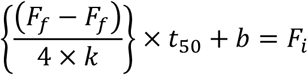

To solve for *b*, at *t_50_* (i.e. the midpoint between *F_f_* and *F_i_*:

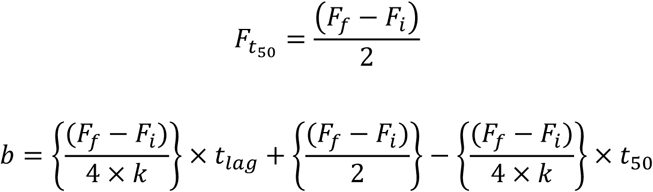

Solving for *t_lag_*:

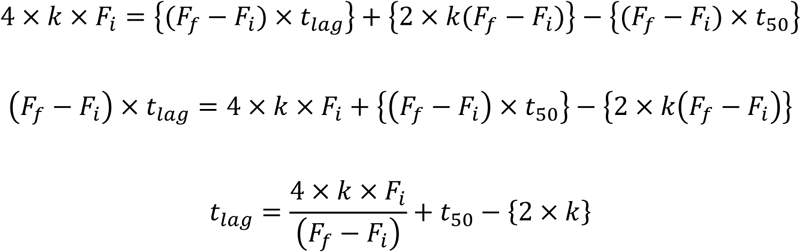

The relative efficacy of each sHsp to inhibit the formation of TDP-43^LCD^ fibrils was determined by calculating the protection conferred by each sHsp according to the difference in maximal ThT fluorescence in the absence and presence of the chaperone using the equation,

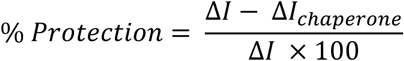

Wherein *ΔI* and *ΔI_chaperone_* correspond to the change in ThT fluorescence of TDP-43^LCD^ in the absence and presence of the sHsp, respectively.

### Kaplan-Meier analysis of TDP-43^LCD^ fibrillation in the presence of sHsps

Kaplan-Meier analysis was performed as previously described (34). We chose to display the data in this way to account for technical replicates in which aggregation was fully suppressed over the course of the experiment. Plots were generated using GraphPad Prism software. To construct the plots from the kinetic data, we used values of *t*_lag_ as the parameter that indicated that an aggregation event had occurred in a particular well on the basis that an increase in ThT fluorescence reflects its binding to a fibril. Each aggregation event was assigned the value “1”. Those wells that did not exhibit any significant increase in ThT were assigned the value “0” to indicate the absence of an aggregation event across the time of the assay. The statistical difference between the Kaplan-Meier plots of each well in comparison to TDP-43^LCD^ alone was determined using the log-rank (Mantel-Cox) algorithm at 95% confidence interval. Hazard ratios were calculated with the log-rank (Mantel-Cox) method using GraphPad Prism 10 software. The hazard ratio for TDP-43^LCD^ in different ratios of sHsp was calculated relative to TDP-43^LCD^ alone. To calculate the mean time to failure, Kaplan-Meier plots were fit with a Boltzmann sigmoidal curve, from which values of *t*_lag_ were derived and assigned as the mean time to failure.

### Transmission electron microscopy

Negative stain transmission electron microscopy (TEM) samples were prepared by applying 5 μL of fibril samples to 400 mesh carbon-coated copper grids (ProSciTech, Queensland, Australia) and incubating on the grid for 30 s. The grids were blotted dry and washed twice with 10 μL filtered water and blotted dry each time before being stained with 2% uranyl acetate for 30 s, blotted dry and allowed to dry for 2 min. Imaging was performed on a T-12 (FEI/Thermo Fisher Scientific) electron microscope.

### Kinetic light scattering assay

To semi-quantify TDP-43^LCD^ phase separation, 20 μL of 40 µM TDP-43^LCD^ was pipetted into a single well of a clear 384-well plate containing 20 μL of 20 mM MES pH 6.0 containing 0 – 1 M NaCl at 100 mM increments, such that the final concentration of protein and NaCl was half of its original concentration. The scattering of light caused by particles including phase separated TDP-43^LCD^ was then measured by optical density (OD) at 340, 400 and 600 nm every 10 s for 30 cycles using a SpectroStar Plate Reader (BMG Labtechnologies, Melbourne, Australia). Assays were conducted using n = 5 biological replicates. At the end of each assay, data for the change in OD were plotted using GraphPad Prism version 10 (Graphpad Software Inc., San Diego, CA). The first 6 data points were fitted with simple linear regression curves to determine the initial rate of change in OD (ΔOD/s). The final data point for each well was plotted to determine the total change in OD (ΔOD) across the time course of the assay.

### Confocal microscopy

To visually assess the extent of TDP-43^LCD^ phase separation, condensates were imaged using differential interference contrast (DIC) confocal microscopy. Condensates were prepared by incubating 50 μL of 40 μM TDP-43^LCD^ for 1 h with an equal volume of NaCl in 20 mM MES (pH 6.0) at concentrations varying from 0 – 1 M in 100 mM increments, such that the protein and NaCl were diluted to half the original concentration. Additionally, condensates were prepared by incubating with an equal volume of 40 μM HspB1^3D^ in 20 mM MES (pH 6.0) or an equal volume of 40 μM HspB5^WT^ in 20 mM MES (pH 6.0) containing 200 mM NaCl. After incubation, 5 μL aliquots were placed on a glass slide with SecureSeal™ adhesive imaging spacers (Thermo Fisher Scientific), which was then sealed with a cover slip. Images were captured using a 63× NA1.2 water immersion objective on a Leica SP5 laser scanning confocal microscope.

### Quantification of phase separated TDP-43^LCD^

To quantify the amount of TDP-43^LCD^ within condensates, samples were incubated at room temperature for 1 h before being centrifuged at 16 300×*g* for 5 min to pellet phase separated material. Supernatants were transferred to a separate tube and their protein concentrations determined using a Nanodrop 2000c spectrophotometer. Data were plotted using GraphPad Prism version 10 (GraphPad Software Inc., San Diego, CA).

### Fluorescent labelling of proteins

TDP-43^LCD^ was dialysed into labelling buffer (20 mM Tris, 500 mM NaCl, 6 M guanidine-HCl, pH 8.0). Following dialysis, 10 mM sodium bicarbonate was added prior to the addition of 8 M Cytiva Cy3 Mono NHS Ester (GEPA13101) (Cy3). Labelled TDP-43^LCD^ was incubated on a rotary mixer overnight at 4°C, after which free Cy3 dye was removed using ZebaSpin^TM^ 7 kDa molecular weight cutoff desalting columns (ThermoFisher). Aliquots of labelled TDP-43^LCD^ were then stored at −80°C. The degree of labelling (DOL) was calculated as follows:

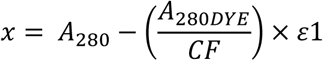

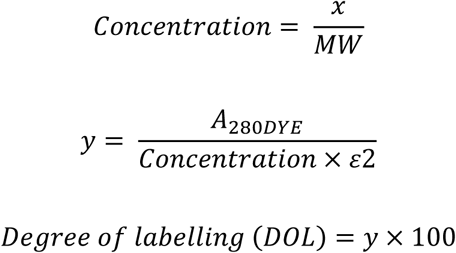

Wherein A_280_ = absorbance of protein measured at 280 nm, A_280DYE_ = absorbance of Cy3 measured at 280 nm, CF = correction factor of Cy3 (0.08), ε1 = extinction coefficient of TDP-43^LCD^ (19480 cm^-1^ M^-1^), MW = molecular weight of TDP-43^LCD^ (17197.43 g mol^-1^), ε2 = extinction coefficient of Cy3.

Fluorescent labelling of HspB1^3D^, HspB5^WT^, HspB1^ACD^, and HspB5^ACD^ was performed as above but using Cy5 Mono NHS Ester (GEPA15101) instead of Cy3. The labelling efficiencies of proteins were as follows: TDP-43^LCD^ (26%), HspB1^3D^ (130%), HspB1^ACD^ (104%), HspB5^WT^ (30%), and HspB5^ACD^ (5.5%).

### Quantification of partitioning

To calculate the free energy of partitioning (ΔG_partition_) within a condensate, confocal fluorescence images of fluorescently labelled TDP-43^LCD^ and HspB1^3D^ or HspB5^WT^ were analysed using ImageJ to determine the mean pixel intensity within a droplet compared to the background solution intensity. Partitioning coefficients were calculated as follows:

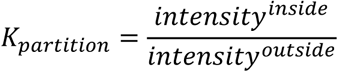

The free energy of partition was calculated as follows:

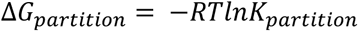

The average ΔG_partition_ was calculated for each protein.

### Fluorescence recovery after photobleaching

To evaluate the effect of HspB1^3D^ and HspB5^WT^ upon the dynamic properties of TDP-43^LCD^, condensates were evaluated via fluorescence recovery after photobleaching (FRAP). Condensates were prepared as per section 2.2.9 with the addition of 1% Cy3-labelled TDP-43^LCD^. FRAP was performed using a Leica SP5 laser scanning confocal microscope equipped with a 63×/1.2 NA water immersion objective at 70% laser power. Bleaching was conducted at 100% laser transmission using the 488 nm line, while 6% laser transmission was using for imaging. PMT detectors were used and calibrated using digital gain to prevent signal oversaturation and 16-bit images were obtained to improve signal dynamic range. Fluorescence recovery was monitored for ∼90 s at intervals of ∼0.171 s.

The fluorescence intensities for the bleached region, reference region (whole condensate) and background region (outside the condensate) were measured and analysed using the program-based tool, EasyFRAP (56). Data underwent full-scale normalisation and mobile fraction and half-time to recovery were derived from these curves manually. At least 5 image series were analysed per sample to calculate the mean and standard deviation (SD). The mean mobile fraction and half-time to recovery for each condition were plotted using Prism version 10 (GraphPad). Data is representative of three independent biological replicates.

### Two-colour coincidence detection (TCCD)

Two-colour coincidence detection (TCCD) was performed using glass bottom 8-well chamber slides (Ibidi, Munich, Germany). Briefly, glass-bottomed chamber slides were blocked for 24 h in blocking buffer (10% (v/v) FCS, 2% (w/v) BSA, 0.1% (v/v) Triton X-100, 1X PBS) before being washed with PBS. Following equilibration to 25°C, Cy3-labelled TDP-43^LCD^ or Cy5-labelled HspB1^3D^, HspB1^ACD^, HspB5^WT^, or HspB5^ACD^ were diluted to 200 nM in 20 mM MES (pH 6.0) immediately prior to imaging. Two-colour coincidence detection (TCCD) was performed using a Leica SP8 FALCON confocal microscope (Leica, Wetzlar, Germany) using an 86×/1.20 water immersion objective using the white light laser (WLL) at 85% output and at a pulse rate of 80 MHz. Cy3-labelled TDP-43^LCD^ was detected using an excitation/emission wavelength of 561/571 nm, while Cy5-labelled chaperones were detected using an excitation/emission wavelength of 647/657 nm. Data were collected using the fluorescence correlation spectroscopy wizard for a period of 10 s for each acquisition, with 25 technical replicates obtained for each experimental replicate (n = 3). Sulforhodamine-B (SRB) was used for dye calibration due to its known diffusion coefficient of 420 µm^2^/s. Purified GFP was used as a non-chaperone control and was detected using an excitation/emission wavelength of 488/498 nm.

### TCCD data analysis

TCCD data was analysed using custom written Python scripts (deposited at Zenodo: DOI: 10.5281/zenodo.138587777). Briefly, the background noise within the fluorescence channels for each treatment were determined by fitting the photo count rate (PCF) to a normal distribution. From this, the peak centre (P_center_) and full width at half-maximum (FWHM) were calculated. For each technical replicate, a peak finding algorithm was applied and only those peaks with a PCF higher than the noise threshold (indicative of large oligomeric species) were identified. To ensure that slowly diffusing oligomers were identified as single peaks, the minimum time between peaks was set to 40 ms.

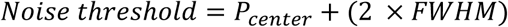

The number of peaks per replicate and their maximum PCFs (a relative indicator of oligomer size) were then determined. For treatments in which two fluorescence channels were measured simultaneously (e.g., TDP-43^LCD^ + GFP), the noise fitting and peak finding was performed on each fluorescence channel separately as described above. To determine whether labelled proteins were forming hetero-oligomers, peaks that were identified in both fluorescence channels within 10 ms of each other were labelled as coincident. Finally, the proportion of TDP-43^LCD^ peaks that were coincident with sHsp or the non-chaperone control (i.e., GFP) was determined:

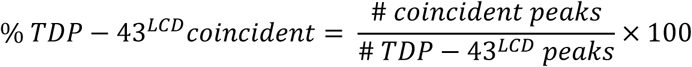

### Statistical analysis

Statistical analysis was performed using Prism software version 10 (GraphPad). Prior to analysis, all data were tested for normal distribution by Shapiro-Wilk test. Data sets with normal distribution were analysed by one-way or two-way ANOVA. Data sets with non-normal distribution were analysed by Kruskal-Wallis test.

