## Supplementary material for "Small heat shock proteins HspB1 and HspB5 differentially alter the condensation and aggregation of the TDP-43 low complexity domain": Supplmentary Files

### Supplementary material description

#### Supplementary Table 1

**Supplementary Table 1. List of bacterial expression vectors.** Plasmids are separated by plasmid backbone and further separated by variant. Sources of plasmids are listed with corresponding references.

| | | |
|---|---|---|
| ***Bacterial expression vector*** | | |
| pJ411 TDP-43_CTD | Contains the gene for bacterial expression of the TDP-43 low complexity domain (LCD) with a 6His-tag and TEV protease site at its N-terminus. The antibiotic resistance is $Kan^r$. | |
| ***Bacterial expression vector*** | | |
| pET3a HspB1 | Contains the gene for bacterial expression of HspB1. The antibiotic resistance is $Amp^r$. | |
| **Variants used in study** | **Source** | **Reference** |
| $HspB1^{WT}$ | Ecroyd group (Wollongong, AU) | (24) |
| $HspB1^{3D}$ | Ecroyd group (Wollongong, AU) | (23) |
| ***Bacterial expression vector*** | | |
| pET24a HspB5 | Contains the gene for bacterial expression of HspB1. The antibiotic resistance is $Kan^r$. | |
| **Variants used in study** | **Source** | **Reference** |
| $HspB5^{WT}$ | Ecroyd group (Wollongong, AU) | (24) |
| ***Bacterial expression vector*** | | |
| pET28-HSP_ACD | Contains the gene for bacterial expression of the α-crystallin domain (ACD) of HspB1 or HspB5. The antibiotic resistance is $Kan^r$. | |
| **Variants used in study** | **Source** | **Reference** |
| $HspB1^{ACD}$ | Laganowsky group (Texas A&M Health Science Center, USA) | (51) |
| $HspB5^{ACD}$ | Laganowsky group (Texas A&M Health Science Center, USA) | (51) |

#### Supplementary Table 2

**Supplementary Table 1. Hazard ratios for TDP-43$^{LCD}$ aggregation in the presence of sHsps.** Hazard ratios were calculated from Kaplan-Meier curves in Figures 2-4 via the log-rank (Mantel-Cox) method to determine how each sHsp altered the likelihood of TDP-43$^{LCD}$ aggregation.

| Molar ratio | HspB1$^{WT}$ | | HspB1$^{3D}$ | | HspB1$^{ACD}$ | | HspB5$^{WT}$ | | HspB5$^{ACD}$ | |
|---|---|---|---|---|---|---|---|---|---|---|
| | Hazard ratio | *P* | Hazard ratio | *P* | Hazard ratio | *P* | Hazard ratio | *P* | Hazard ratio | *P* |
| 5:1 | 0.1985 | <0.0001 | *N/A* | *N/A* | *N/A* | *N/A* | 0.1026 | <0.0001 | *N/A* | *N/A* |
| 1:1 | 0.2057 | <0.0001 | 0.2515 | 0.0002 | 0.4884 | ns | 0.1694 | <0.0001 | 1.003 | ns |
| 1:5 | 0.2124 | <0.0001 | 0.2267 | 0.0002 | *N/A* | *N/A* | 0.1417 | <0.0001 | *N/A* | *N/A* |
| 1:10 | 0.2268 | <0.0001 | 0.2610 | 0.00035 | 1.1690 | ns | 0.1417 | <0.0001 | 1.3800 | ns |
| 1:100 | 0.5383 | ns | 0.2918 | 0.0423 | 1.2110 | ns | 0.2834 | 0.0005 | 3.4980 | 0.0014 |

**Supplementary Figure 1**

TDP-43$^{LCD}$ incubated at 50 μM rapidly became turbid, which was indicative of it having undergone condensation at these concentrations (**Fig. S1a**). The kinetics of TDP-43$^{LCD}$ aggregation at this concentration and at 20 μM were reduced compared to 10 μM TDP-43$^{LCD}$ (**Fig. 1**), indicating that TDP-43$^{LCD}$ condensation alters its aggregation.

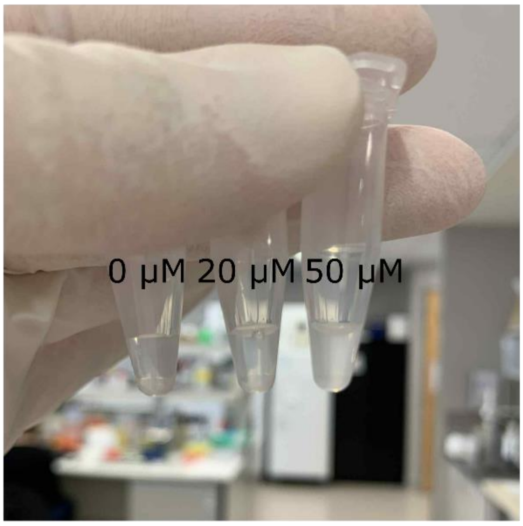


**Supplementary Figure 1.** High concentrations of TDP-43$^{LCD}$ exhibit turbidity indicative of condensation.

#### Supplementary Figure 2

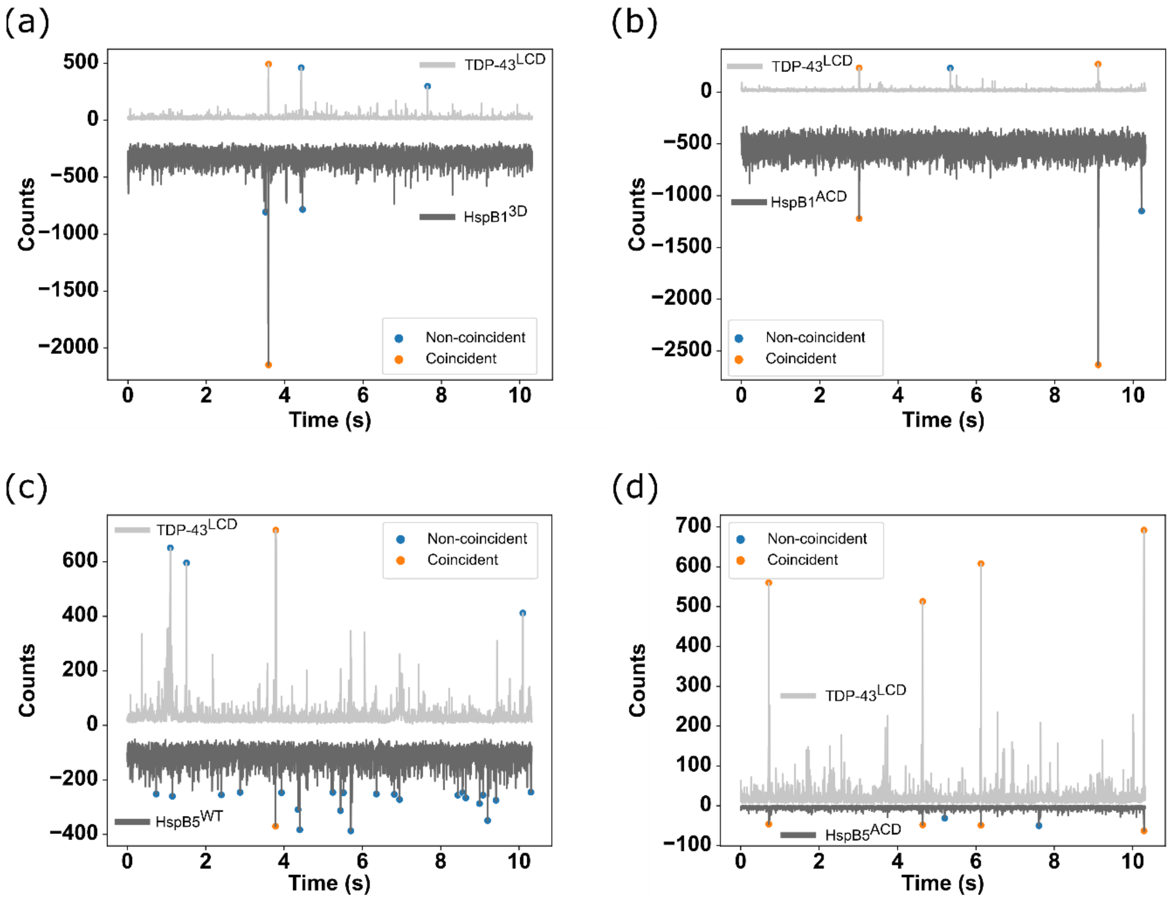


**Supplementary Figure 2.** Two-colour coincidence detection (TCCD) was performed using a Leica TCS SP8 FALCON laser scanning confocal microscope. (a-d) Fluorescence intensity traces of Cy3-TDP-43$^{LCD}$ with (a) HspB1$^{3D}$, (b) HspB1$^{ACD}$, (c) HspB5$^{WT}$, and (d) HspB5$^{ACD}$. TDP-43$^{LCD}$ is indicated in light grey. sHsps are shown in dark grey. Coincident peaks are indicated by orange circles while non-coincident peaks are indicated by blue circles.

#### Supplementary Figure 3

To determine whether partitioning to TDP-43$^{LCD}$ condensates was a specific property of HspB1 and HspB5, TDP-43$^{LCD}$ was incubated with the non-chaperone protein mCherry and 200 mM NaCl. As expected, mCherry was excluded from TDP-43$^{LCD}$ condensates (**Fig. S2a**), indicating that HspB1 and HspB5 localisation to these condensates was specific. Importantly, mCherry remained excluded even after 5 h of incubation (**Fig. S2b**), indicative of minimal diffusion of protein into TDP-43$^{LCD}$ condensates.

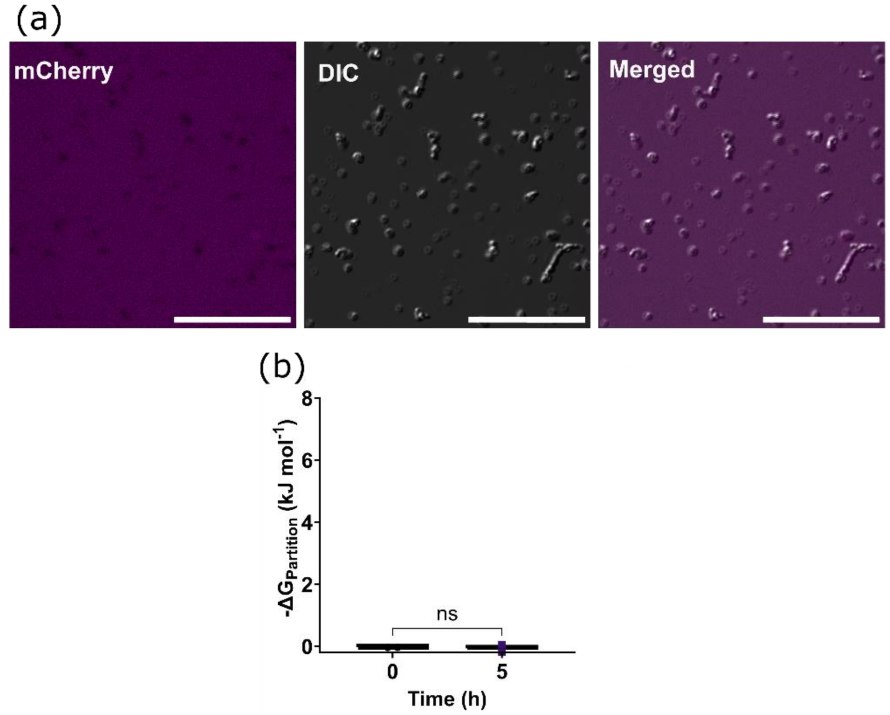


**Supplementary Figure 2.** TDP-43$^{LCD}$ (20 μM) was incubated with mCherry (20 μM) and 200 mM NaCl and imaged using a Leica TCS SP5 laser scanning confocal microscope. Partitioning of mCherry into TDP-43$^{LCD}$ condensates was calculated. (a) Confocal microscopy images of mCherry partitioning. Scale bar represents 20 μm. (b) Partitioning of mCherry to TDP-43$^{LCD}$ condensates.
